# Redox signaling regulates breast cancer metastasis via HIF1α-stimulated EMT dynamics and metabolic reprogramming

**DOI:** 10.1101/2022.08.29.503508

**Authors:** Zuen Ren, Malindrie Dharmaratne, Huizhi Liang, Outhiriaradjou Benard, Miriam Morales-Gallego, Kimita Suyama, Atefeh Taherian Fard, Jessica C. Mar, Michael Prystowsky, Larry Norton, Rachel B. Hazan

## Abstract

Metastasis is orchestrated by phenotypic and metabolic reprogramming underlying tumor aggressiveness. Redox signaling by mammary tumor knockdown (KD) of the antioxidant glutathione peroxidase 2 (GPx2) enhanced metastasis via dynamic changes in epithelial-to-mesenchymal transition. Single cell RNA sequencing (scRNA-seq) of the control and PyMT/GPx2 KD mammary tumor revealed six luminal and one basal/mesenchymal like (cluster 3) subpopulations. Remarkably, GPx2 KD enhanced the size and basal/mesenchymal gene signature of cluster 3 as well as induced epithelial/mesenchymal (E/M) clusters which expressed markers of oxidative phosphorylation and glycolysis, indicative of hybrid metabolism. These data were validated in human breast cancer xenografts and were supported by pseudotime cell trajectory analysis. Moreover, the E/M and M states were both attenuated by GPx2 gain of function or HIF1α inhibition, leading to metastasis suppression. Collectively, these results demonstrate that redox/HIF1α signaling promotes mesenchymal gene expression, resulting in E/M clusters and a mesenchymal root subpopulation, driving phenotypic and metabolic heterogeneity underlying metastasis.

**Significance:** By leveraging single cell RNA analysis, we were able to demonstrate that redox signaling by GPx2 loss in mammary tumors results in HIF1α signaling, which promotes partial and full EMT conversions, represented by distinct tumor cell subpopulations, which in turn express hybrid and binary metabolic states. These data underscore a phenotypic and metabolic co-adaptation in cancer, arguing in favor of the GPx2-HIF1α axis as a therapeutic platform for targeting tumor cell metastasis.

## Introduction

Metastasis is a systemic disease that is often incurable with currently available therapies (1). It is believed that tumor heterogeneity drives metastasis by selecting for tumor cell subpopulations that are endowed with EMT and stemness (2, 3). Cancer cell hyper-proliferation leads to cell crowding, resulting in depletion of nutrients and oxygen, leading to hypoxia. Tumors tend to use aerobic glycolysis or the Warburg effect to build energy and biomass. However, tumors may also only use mitochondrial oxidative phosphorylation, which increases ROS production and hence HIF1*α* stabilization, leading to tumor stemness and chemoresistance (4, 5). HIF1*α* is a master regulator of key cellular processes, involving cell proliferation, survival, migration, metabolism, and importantly EMT, which drives stemness and metastasis. HIF1*α* is known to play a pivotal role in EMT via transcriptional activation of EMT factors like *Snai1/2* and *Zeb1*, which suppress epithelial differentiation via repression of epithelial genes, especially E-cadherin (6, 7).

Our recent work highlights the profound effects of redox signaling by knockdown of GPx2 in mammary tumors, resulting in ROS/HIF1α signaling, causing vascular malfunction and hypoxia, leading to metabolic heterogeneity (8). In fact, GPx2 downregulation is a physiologically relevant change in breast cancer. Analysis of a database of 1,809 patients showed an association between low GPx2 mRNA in luminal B, HER2-enriched, basal-like breast tumors and poor patient survival (8). Examination of similarly-sized control and GPx2 KD tumors confirmed that GPx2 KD actively promotes metastasis, thus ruling out hypoxia caused by tumor bulkiness as the main impetus for advancing malignancy (8). The mechanism underlying metastatic progression by GPx2 KD remained however unclear. EMT is a pivotal process which transforms epithelial carcinoma cells into mesenchymal ones, resulting in acquisition of stem and metastatic properties (9-12). However, recent studies have shifted our view on EMT, indicating it is a dynamic rather than a binary process, that gives rise to intermediate epithelial/mesenchymal (E/M), or even a continuum of E/M states, further exacerbating phenotypic heterogeneity and metastasis (13-17). Interestingly, the E/M hybrid state is thought to be represented by a minor subset of carcinoma cells in basal breast cancers, that is able of initiating tumors and promoting metastasis (14-18). In light of these findings, we sought to investigate whether GPx2 KD promotes EMT dynamics at the single cell level, and unravel unique tumor cell subpopulations and/or oncogenic drivers governing metastasis.

Using scRNAseq of the PyMT/GPx2 KD *versus* the PyMT control tumor model, we identified an overlap in cell clustering based on cell type and cell state involving six luminal-like clusters and one basal/mesenchymal-like cluster (cluster 3). Cluster 3 showed striking upregulation of basal/mesenchymal genes and downregulation of luminal genes in cluster 3 relative to all luminal clusters. Remarkably, GPx2 KD increased the pool size and exacerbated the mesenchymal signature of cluster 3. Moreover, GPx2 KD induced striking *de novo* expression of mesenchymal genes in luminal clusters, thus generating an extreme basal/mesenchymal-like cluster (cluster 3) and several epithelial/mesenchymal (E/M) clusters. These findings were consistent with high incidence of areas expressing E/M or M genes in GPx2 KD tumors relative to controls. These findings were corroborated by pseudotime cell trajectory analysis, underscoring cluster 3 as a root or stem-like subpopulation. Importantly, E/M tumors expressed a hybrid metabolic gene signature, involving phosphorylated-AMPK and GLUT1, indicative of OXPHOS and glycolysis (8). All of these effects were interestingly reversed by GPx2 overexpression or HIF1α inhibition, resulting in dramatic suppression of spontaneous metastasis. These data underscore a dramatic effect of redox regulation of HIF1α signaling on EMT dynamics, resulting in phenotypic and metabolic co-adaptation that may be leveraged for therapeutic intervention in metastatic breast cancer.

## Results

### The identification of a basal/mesenchymal-like tumor cell subpopulation at the single cell resolution

Previously, we showed that GPx2 KD in the PyMT model dramatically enhanced spontaneous lung metastasis (8); however, the mechanism remained unknown. EMT is a hallmark of cancer metastasis and a process whereby tumors dynamically transform epithelial cells into mesenchymal-like cells (19, 20). During EMT, tumor cells gain motility, invasiveness and stemness properties which drive metastasis, recurrence or chemoresistance (9-12). scRNAseq of FACS-sorted GFP-labelled cells from one PyMT1/GPx2 KD and one PyMT1/control mammary tumor unraveled seven epithelial tumor cell subpopulations (clusters) that were shared by the GPx2 KD and control tumor (8). Of these, six were luminal-like (cluster 0, 1, 2, 4, 5, 6) and one (cluster 3) was basal/mesenchymal-like. Cluster identification was based on manually supervised annotation using MSIGDB hallmark genes indicative of cell type (basal, luminal) or cell state (proliferation status) (**Fig. 1A**). Cluster 3 exhibited a classical EMT/stem-like gene signature involving upregulation of basal/mesenchymal genes (*Vim, Krt14, Klf4, Jag1, Notch1, Aldh2, Itgb4, Twist, Sparc, S100a4*), and downregulation of luminal/epithelial genes (*Krt8/18, Epcam, Cdh1, Cldn3/7*) (**Fig. 1A**). Of note, expression of the stemness genes *Klf4, Jag1, Notch1, Aldh2* in cluster 3, implies it may harbors stemness activity. By contrast, luminal cluster 2 was enriched in a proliferative gene signature (*Mki67, Ccnb2, Cdk1, Ccnb1 and Ccnd1*) (**Fig. 1A**), indicating it might contain cycling progenitor cells, giving rise to the bulk of the tumor. In addition, we identified two non-epithelial clusters (clusters 7 and 8), expressing macrophage and fibroblast like signatures which may be tumor-associated stromal cells that were co-sorted with tumor cells (**Fig. 1A**).

**Fig 1.**
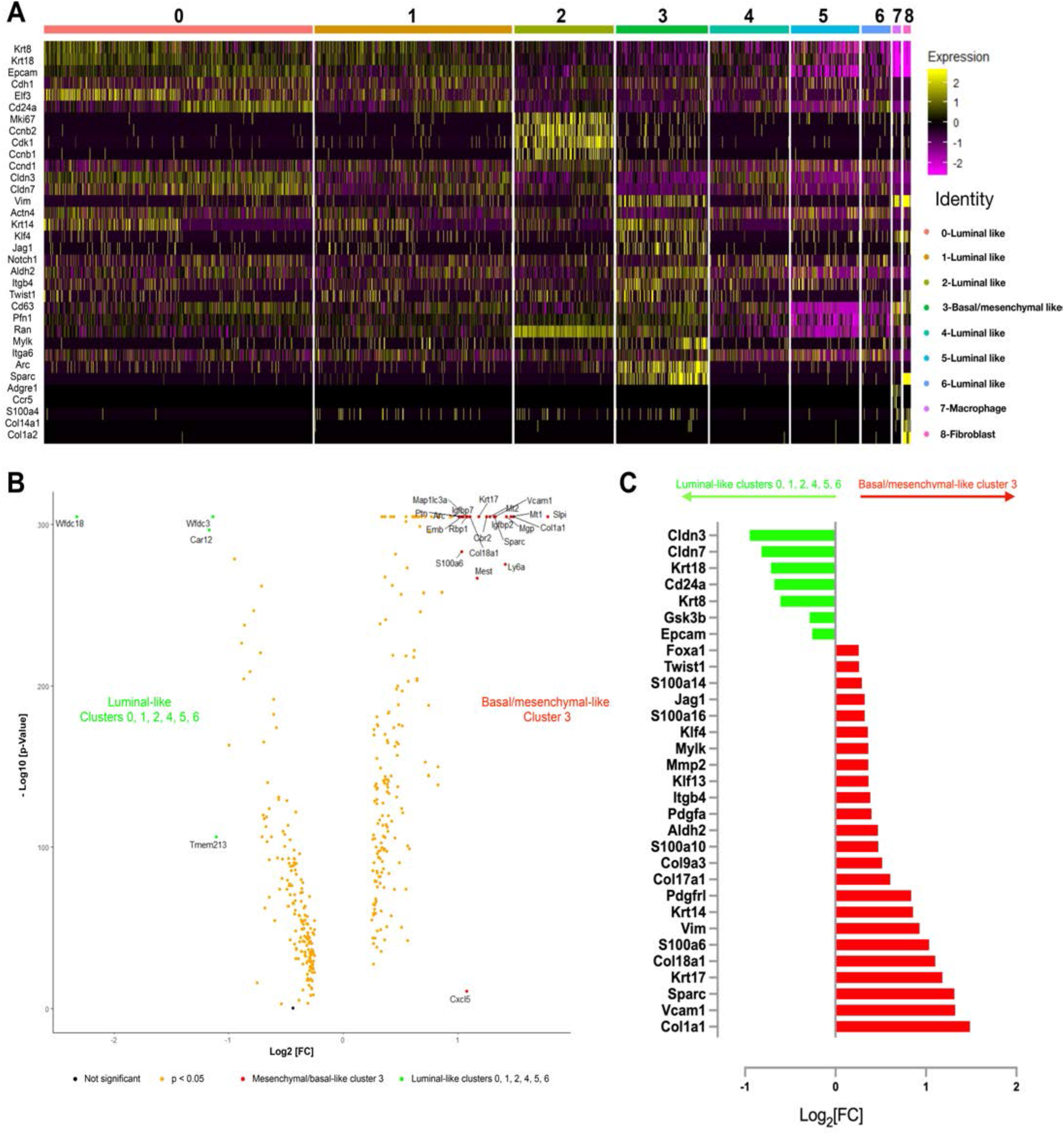
scRNAseq reveals a subpopulation of cells in PyMT mammary tumor model enriched in upregulation of basal/mesenchymal genes. **(A)** Heatmap shows manually supervised-automated clustering of individual interested marker genes. **(B)** Volcano plot shows the differentially expressed genes between cluster 3 (cell number: 1666) and clusters 0, 1, 2, 4, 5, 6 (cell number: 13433). X-axis is a Log transformation of mRNA fold change at base 2. Y-axis is a negative Log transformation of p-Value at base 10. Yellow dot indicates significantly expressed genes (p < 0.05); Green dot highlights top genes with log2[FC] less than -1 and -log10 [p-Value] greater than 100 that are significantly upregulated in luminal-like clusters 0, 1, 2, 4, 5, 6. Red dot highlights top genes with log2 [FC] greater than 1 and -log10 [p-Value] greater than 100 that are significantly upregulated in basal/mesenchymal-like cluster 3. (**C**) Bar graph shows log2[FC] values for the differentially and significantly expressed genes-associated with EMT and stemness in cluster 3 vs. clusters 0, 1, 2, 4, 5, 6 (p < 0.05).

To further confirm the clustering data obtained by manual annotation, we unbiasedly analyzed the differential gene expression between cluster 3 and all luminal-like clusters combined (cluster 0, 1, 2, 4, 5, 6), resulting in volcano plot illustrating the highly basal/mesenchymal-like gene signature of cluster 3. This was indicated by striking upregulation of the basal/mesenchymal *Krt17, Col1a1, Vcam1, Sparc, Col18a1, S100a6* genes in cluster 3 relative to all luminal clusters (**Fig. 1B**). Of note, Krt17 is a basal keratin that is often expressed in mesenchymal TNBCs. Further, analysis of EMT- and stemness-associated genes in cluster 3 *versus* all luminal clusters, revealed an over-representation of basal/mesenchymal genes (*Vim, Krt14, Krt17, Pdgf, Sparc, Arc, Mest, S100a6, Vcam1, Col1a1, Col18a1*) and EMT/stemness genes (*Foxa1, Twist1, Jagged1, Klf4, Itgb4*) in cluster 3 (**Fig. 1C**). These effects were accompanied by downregulation of luminal lineage genes (*Cldn3, Cldn7, CD24, Krt18, Krt8, Epcam*) in cluster 3, highlighting that this subpopulation resides in a mesenchymal state (**Fig. 1C**). In further support of this notion, Ingenuity Pathway Analysis (IPA) of cluster 3 relative to all luminal clusters, showed enrichment in EMT pathways and motility regulatory processes involving Integrin, Actin cytoskeleton and RhoA signaling (**Fig. S1A**). In support of stemness activity, cluster 3 was enriched in IL-6 signaling and NRF2 anti-oxidant response, which might prevent CSC differentiation by ROS (21-23) (**Fig. S1A**). Together, these results support the notion of cluster 3 resides in a mesenchymal state endowed with stemness traits.

### GPx2 KD in the PyMT tumor potentiates EMT process and induces de novo mesenchymalization of luminal clusters

Our previous data implied that GPx2 KD potentiates a binary E to M switch to enhance metastasis (8). Thus, we sought to compare the various clusters in the GPx2 KD versus control mammary tumor in UMAPs (**Fig. 2A**). Interestingly, analysis of the population size of each of the clusters unraveled a significant increase in cell number in cluster 3 in the GPx2 KD tumor relative to control tumor: 983 (12.65%) vs 683 (8.96%), (Pearson’s Chi-square test, p-value = 2.2e-16; two-sample test for equality of proportions with continuity correction data c (983, 683) out of c (7768, 7621), p-value = 1.03e-13) (**Fig. 2B-C**). Conversely, luminal clusters 4 and 5 were reduced in size in the GPx2 KD tumor relative to control tumor (from 10.75% to 7.87%, and 9.38% to 6.69%, respectively) (**Fig. 2B-C**). Cluster 2 was increased from 10.23% in control tumor to 13.16% in the GPx2 KD tumor, consistent with a proliferative gene signature driving tumor growth (**Fig. 2C**) (8). These results implied that GPx2 KD expands cluster 3 while contracting clusters 4 and 5. Namely, variations in subpopulation size may reflect changes in cluster identity, likely due to tumor cell “mesenchymalization” by EMT activation by GPx2 KD, thus converting cells in luminal cluster 4 or 5 into cluster 3-like cells. Hence, we used our scRNAseq data to examine the effect of GPx2 KD on the expression of classical basal/mesenchymal and luminal genes in the various clusters. Indeed, mRNA expression analysis by feature plot showed that GPx2 KD increased the expression of basal keratins (*Krt5, Krt14, Krt17*) and mesenchymal genes (*Vim, Twist1/2, Cdh2* (N-cadherin)) (**Fig. 2D-E**), while reducing *Epcam* and *Claudin 7* in most clusters; with *Claudin 7* more significantly reduced in cluster 3 (**Fig. 2F**). Together, these results support that GPx2 KD dynamically shift cells in cluster 4 and 5 from a luminal to an M state, thereby augmenting the size of cluster 3.

**Fig. 2.**
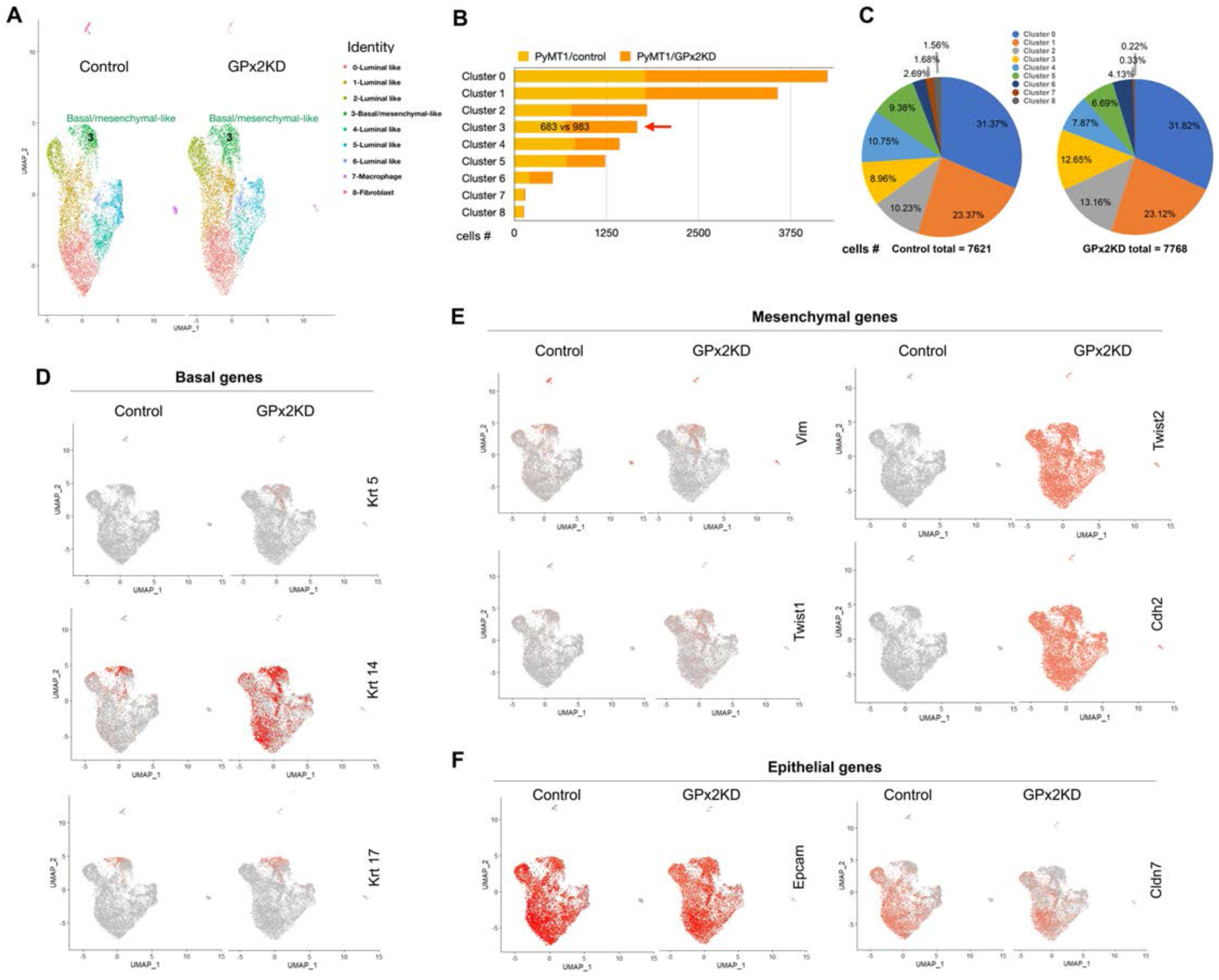
GPx2 KD potentiates mesenchymal cluster 3 and induces mesenchymal gene expression in luminal clusters. (**A**) UMAP projection of comprehensively integrated clustering results from one PyMT1 control and one PyMT1/GPx2 KD tumor revealed six luminal-like clusters (cluster 0, 1, 2, 4, 5, 6), one basal/mesenchymal-like cluster (cluster 3), and two are non-epithelial clusters (cluster 7 and 8). (**B**) Bar graphs display the number of cells in each cluster in GPx2 KD versus control. (**C**) Pie graphs display the percentage of cells in each cluster in GPx2 KD versus control. Pearson’s Chi-square test, p-value = 2.2e-16; two-sample test for equality of proportions with continuity correction, data c(983, 683) out of c(7768, 7621), p-value = 1.03e-13. (**D-F**) Feature plots in low-dimensional space showing expression of classical basal marker genes (*Krt5, Krt14, Krt17*) (D), mesenchymal marker genes (*Vim, Twist1, Twist2, Cdh2*) (E), and epithelial marker genes (*Epcam, Cldn7*) (F) in the GPx2 KD relative to control tumor.

Remarkably, GPx2 KD stimulated striking *de novo* induction of basal *Krt14* and mesenchymal *Twist2* and *Cdh2* genes (**Fig. 2D-E**), in cells expressing epithelial *Epcam, Krt8, Kr18*, and *Cldn3* (**Fig. 2F** and **Fig. S2A**). These data underscored the dramatic effect of GPx2 KD on EMT dynamics, converting luminal clusters into a hybrid E/M like state, thus generating cells with partial EMT traits. Interestingly, the E/M phenotype was shown to be conducive of metastasis in many cancers (14, 16, 17), which was in line with the upregulation of invasion-promoting and pro-angiogenic *Mmp2* in several clusters, including cluster 3 (**Fig. S2B**) (24). By contrast, *Snai1/2* mRNA was unchanged (**Fig. S2B**); however SNAI1 protein was markedly upregulated in GPx2 KD tumors (shown later in Fig. 3A), likely due to reduced *Gsk3β* level and thus activity (**Fig. S2C**), resulting in SNAI1 hypo-phosphorylation, preventing ubiquitin-mediated degradation (25). In fact, Wnt signaling (measured by TOP/FOP Flash TCF1 reporter activity) was increased in GPx2KD cells grown under hypoxia (**Fig. S2D**), thus providing a link between GPx2KD, hypoxia, Wnt-induced GSK3*β* inactivation and SNAI1 protein stabilization. Finally, in support of stemness potential, *CD49f* mRNA was increased while *CD24* or *Epcam* mRNA was decreased in most GPx2 KD clusters (**Fig. S2E, Fig. 2F**), which is consistent with CD49f^high^/CD24^low^ or CD49f^high^/Epcam^low^ as CSC markers (26, 27). Jagged1 (Jag1) mRNA was upregulated by GPx2 KD in cluster 3, which may activate stem signals by Notch1 (28) (**Fig. S2E**). Collectively, these data demonstrate that GPx2 KD promotes mesenchymal gene expression, thus generating a continuum of EMT states involving E/M and M tumor cell subpopulations, driving stemness, invasiveness, and hence metastasis. These findings were in keeping with the view that EMT is not merely a binary switch but a dynamic process creating a continuum of phenotypic states that contribute to tumor heterogeneity and aggressiveness (14-18).

**Fig. 3.**
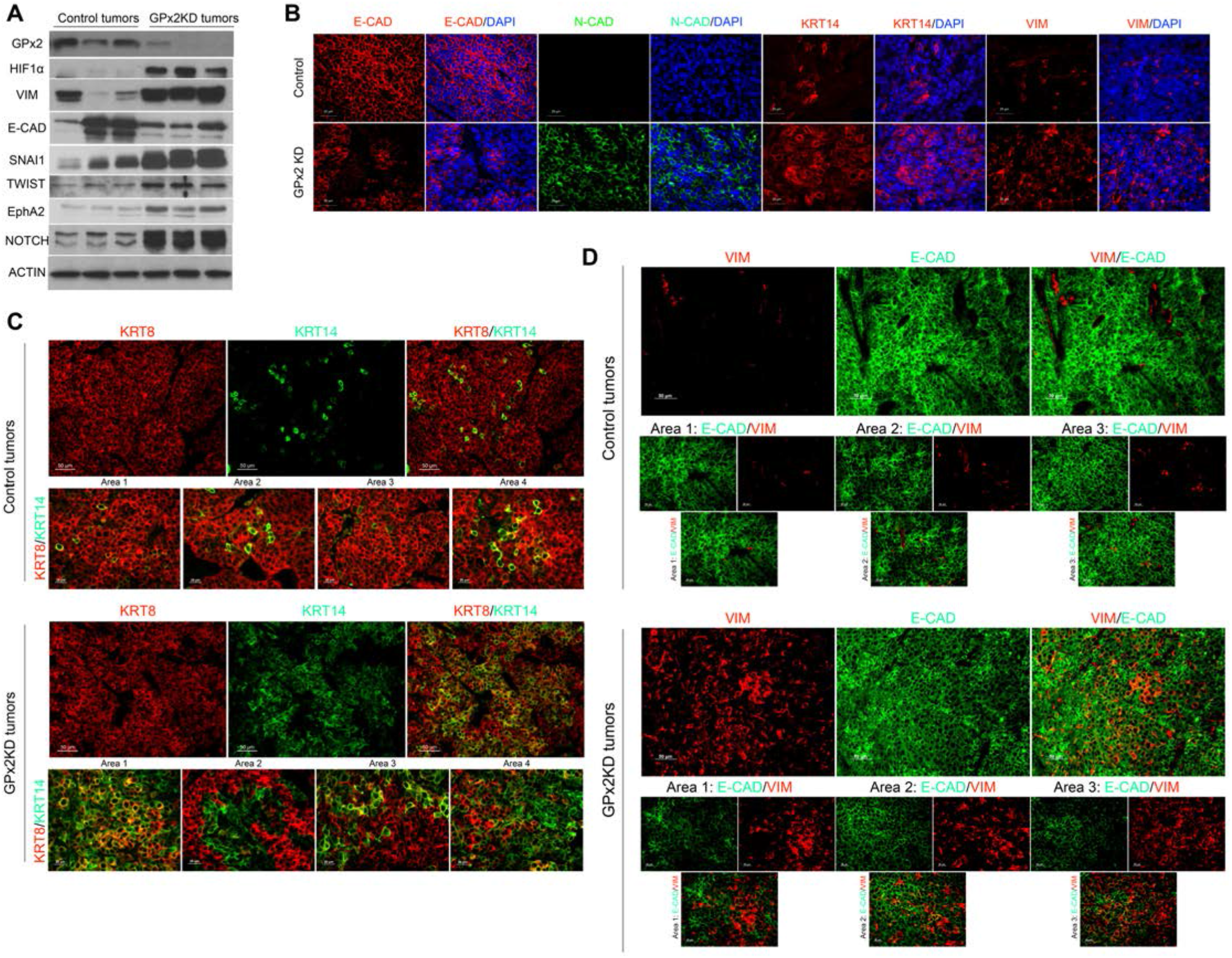
GPx2 loss promotes EMT dynamics *in vivo*. (**A**) Western blots comparing lysates from PyMT1/GPx2 KD to control PyMT1 mammary tumors, derived from 3 independent mice, show changes in expression level of GPx2, HIF1α, VIM, E-CAD, SNAI1, TWIST, EphA2, NOTCH1, relative to ACTIN. (**B**) Immunofluorescent staining of tumors for E-CAD, N-CAD, KRT14, and VIM in PyMT1 control tumors and PyMT1/GPx2 KD tumors from 5 mice (2 tumors per mouse). Three random sections from each tumor were immunostained. Representative images are shown. (**C-D**) Immunofluorescent staining for KRT8 and KRT14 (C), VIM and E-CAD (D) in PyMT1 control tumors and PyMT1/GPx2 KD tumors (n=10) from 5 mice; Representative images illustrate different EMT states in different tumor areas; upper panels, 20X magnification; lower panels, 40X magnification

### Cell trajectory analysis highlights the effect of GPx2 KD on dynamic EMT states between epithelial and mesenchymal cells

These data implied that the M state of cluster 3 in the GPx2 KD tumor represents a highly undifferentiated state conducive of stemness. However, E/M cancer cells were shown to have higher tumor initiating and metastatic potential than M cancer cells (14-18, 29). To examine the transcriptional dynamics linking epithelial to mesenchymal subpopulations in the tumor, we leveraged our single cell analysis to investigate the temporal evolution of EMT in a continuum across all clusters. We employed a computational method to define cell trajectories of epithelial and mesenchymal lineage states across all clusters using Monocle 3 (30, 31). We found that the main trajectory was consistent with our pre-set clustering results (**Fig. S3A-B**). Of note, cell trajectory and UMAP clustering projections were clearly indicative of phenotypic conversions between M and E states, underscoring a rapport between mesenchymal and luminal cell lineages (**Fig. S3B**). To determine how mesenchymal cluster 3 relates to the rest of the luminal clusters, we placed cluster 3 at the root of the trajectory representing the most mesenchymal transcriptional region (**Fig. S3B**). Consistently, cluster 3 exhibited the highest canonical mesenchymal and the lowest canonical luminal gene expression. These results were confirmed by monitoring the expression of classical epithelial or mesenchymal marker genes that rise and fall at different rates across pseudotime distance between the different clusters. For example, *Krt14*, a known basal cell marker, was highly expressed in cluster 3 with greatest pseudotime distance from the utmost luminal cluster (cluster 0) (**Fig. S3C**). Interestingly, *Krt14* displayed a slightly variable pattern in transition state across the clusters, more specifically falling away from cluster 3, rising again in cluster 5, and declining to lowest level in cluster 0 (**Fig. S3C**), pointing to cluster 5 as a one of the earliest luminal clusters transiting into an E/M state. By contrast, luminal *Krt18* expression was lowest in cluster 3, slowly rose in luminal clusters and plateaued in luminal cluster 0 (**Fig. S3D**). Similarly, *Epcam* which was lowest in cluster 3, gradually rose and plateaued across the luminal clusters (**Fig. S3E**). Conversely, the expression of the ultra-mesenchymal gene *S100a6*, and to a lesser extent also that of *Col1a1*, was highest at the root mesenchymal cluster 3, but gradually declined to lowest level across the luminal clusters (**Fig. S3F-G**). These results suggested that GPx2 KD promotes dynamic changes in E to M transitions, generating tumor cell subpopulations expressing a continuum of EMT states. The pseudotime trajectory implies that cluster 3 may contain cells residing at the apex of the CSC hierarchy, which give rise to luminal progenitors while maintaining an ancestral cancer stem cell pool.

### GPx2 knockdown promotes EMT dynamics in tumors in vivo

To validate these findings *in vivo* in the tumor context, we examined the relative expression of epithelial and mesenchymal markers in PyMT/GPx2 KD *versus* control tumors. Immunoblotting of mammary tumors revealed marked increases in HIF1α, a known transcriptional activator of mesenchymal genes (6, 32), hence consistent with the upregulation of VIM, SNAI1, TWIST and downregulation of E-CAD in GPx2 KD tumors (**Fig. 3A**). Since immunoblotting of random tumor chunks may not reflect the heterogeneous distribution of EMT markers in the tumor mass, we immunostained tumor sections for luminal and mesenchymal markers to obtain an overview of the tumor landscape. Indeed, KRT14, N-CAD and VIM proteins were all upregulated whereas E-CAD was downregulated in GPx2 KD tumors (**Fig. 3B**). In addition, EphA2 and NOTCH1, which are implicated in EMT and stemness, were both upregulated in GPx2 KD tumors (**Fig. 3A**) (33, 34). NOTCH1 may also play a role in abnormal angiogenesis leading to hypoxia in GPx2KD tumors (8), as well as in stemness (8). In sum, these data substantiate our scRNA-seq, pointing to the effects of GPx2 KD on EMT and stemness, likely via HIF1α signaling driving the onset of E/M and M tumor cell subpopulations.

To validate these data *in vivo*, we stained tumors for basal KRT14 and luminal KRT8, which were co-expressed in luminal clusters. Indeed, GPx2 KD tumors exhibited hybrid areas with striking co-localization of KRT14 and KRT8 as compared to control tumors which were predominantly KRT8 positive (**Fig. 3C**; Areas 1, 3, 4). Of note, some carcinoma areas in GPx2 KD tumors were either KRT14 or KRT8 positive (**Fig. 3C;** Area 2). Together, these data reflect an EMT spectrum made of luminal, luminal/basal, and basal like clusters. Similar data were obtained when tumors were stained for E-CAD and VIM. Namely, while control tumors were E-CAD^high^ and VIM^low^, GPx2 KD tumors contained areas (especially Area 3, but also Area 2) that were E-CAD^high^ and VIM^high^ (**Fig. 3D**). However, some cell clusters were either E-CAD^high^ or VIM^high^ (especially Area1), pointing to mutually exclusive epithelial or mesenchymal differentiation (**Fig. 3D**). In sum, these data underscore the notion that GPx2 KD promotes tumor heterogeneity, resulting in distinct tumor zones residing in E, M, or E/M state.

Further, to test the contribution of the E/M hybrid phenotype to metastasis, we studied EMT in cell lines that were derived from the same parental PyMT mammary tumor, and were of low (PyMT1) or high (PyMT2) metastatic potential (8). Of note, PyMT2 were GPx2^low^/ROS^high^ with a fibroblastic morphology, whereas PyMT1 cells were GPx2^high^/ROS^low^ with an epitheloid cell shape (**Fig. S4A**) (8). Consistent with mesenchymal differentiation, PyMT2 cells overexpressed N-CAD, SNAIL, SLUG, and VIM protein relative to PyMT1 cells, whereas E-CAD was unchanged (**Fig. S4B**), indicative of an E/M phenotype. Mammary tumors generated by these cell lines showed dramatic upregulation of mesenchymal genes in PyMT2 relative to PyMT1 controls (**Fig. S4C**). PyMT2 tumors over-expressed KRT14, N-CAD, SNAI1, SLUG, VIM and under-expressed E-CAD relative to PyMT1 tumors (**Fig. S4C**), indicative of a binary E to M switch. However, immunostaining revealed that PyMT2 tumors were dense in areas co-expressing KRT8 and KRT14 (Areas 2 and 3) relative to PyMT1 tumor, except for Area 1 showing regions with non-overlapping expression of these markers (**Fig. S4D**). Further, PyMT2 tumors showed areas co-expressing E-CAD and VIM (especially Area 2 and 3) as compared to none in PyMT1 tumors (**Fig. S4E**). Hence, enrichment of E/M clusters in highly-metastatic PyMT2 relative to lowly-metastatic PyMT1 tumors supports a link between the E/M hybrid phenotype and metastasis.

### GPx2 overexpression (OE) in PyMT2 cells suppresses EMT and metastasis

GPx2 overexpression in PyMT2 cells suppresses invasion and spontaneous lung metastasis (8). Consistently, GPx2 OE caused a dramatic shift from a mesenchymal to an epithelial morphology, thus reversing EMT (**Fig. S5A**). In fact, PyMT2/GPx2 OE cells downregulated N-CAD, SNAI1 and SLUG protein expression relative to PyMT2 control cells (**Fig. S5B**). *In vivo*, GPx2 OE tumors increased E-CAD and decreased N-CAD, KRT14, and VIM protein relative to controls (**Fig. S5C**), implying GPx2 promotes mesenchymal to epithelial transition (MET). Interestingly, compared to PyMT2 tumors which harbored KRT8/KRT14 or E-CAD/VIM double positive areas, GPx2 OE tumors were predominantly KRT8 or E-CAD positive, implying an E/M to E state transition (**Fig. S5D-E**). Of note, some VIM^+^ elongated cells in control tumors may be carcinoma-associated fibroblasts (**Fig. S5E**). These data underscored that GPx2 OE in PyMT2 tumors suppresses the E/M or M phenotypes, resulting in luminal/epithelial non-metastatic tumors (8).

### GPx2 de-repression in PyMT2 cells recapitulates the effect of GPx2 OE on EMT and metastasis

The absence of GPx2 in PyMT2 cells might be due to transcriptional or epigenetic silencing. We used the CRISPR/dCas9 system to endogenously induce GPx2 in PyMT2 cells and test similarities with GPx2 OE. We fused dead-Cas9 (dCas9) to the transcriptional activator VP64-p65-Rta (VPR) and customized GPx2 single-guide RNAs (sgRNAs) (GPx2-gRNA1 and GPx2-gRNA2) that direct dCas9-VPR to the GPx2 proximal promoter region for de-repression (**Fig. 4A**). As control, we used a dCas9-VPR without sgRNA. sgRNA2 was the most effective in inducing GPx2 protein, quenching ROS and inducing an epitheloid morphology as compared to control sgRNA (**Fig. 4B-D**). In vivo, PyMT2/ gRNA2 cells were markedly reduced in tumorigenic or metastatic ability as compared to control cells (**Fig. 4E-F**). Consistent with an effect of GPx2 on MET, PyMT2/gRNA2 tumors overexpressed E-CAD and downregulated N-CAD, VIM, and KRT14 (**Fig. 4G-H**). Moreover, PyMT2/gRNA2 tumors were devoid of KRT8/KRT14 or E-CAD/VIM co-expressing areas as compared to controls, resulting in KRT8/E-CAD positive tumors (**Fig. 4I-J**). These data underscored that GPx2 induction, as GPx2 OE, suppressed E/M and M phenotypes, resulting in epithelial differentiation suppressing malignancy.

**Fig. 4.**
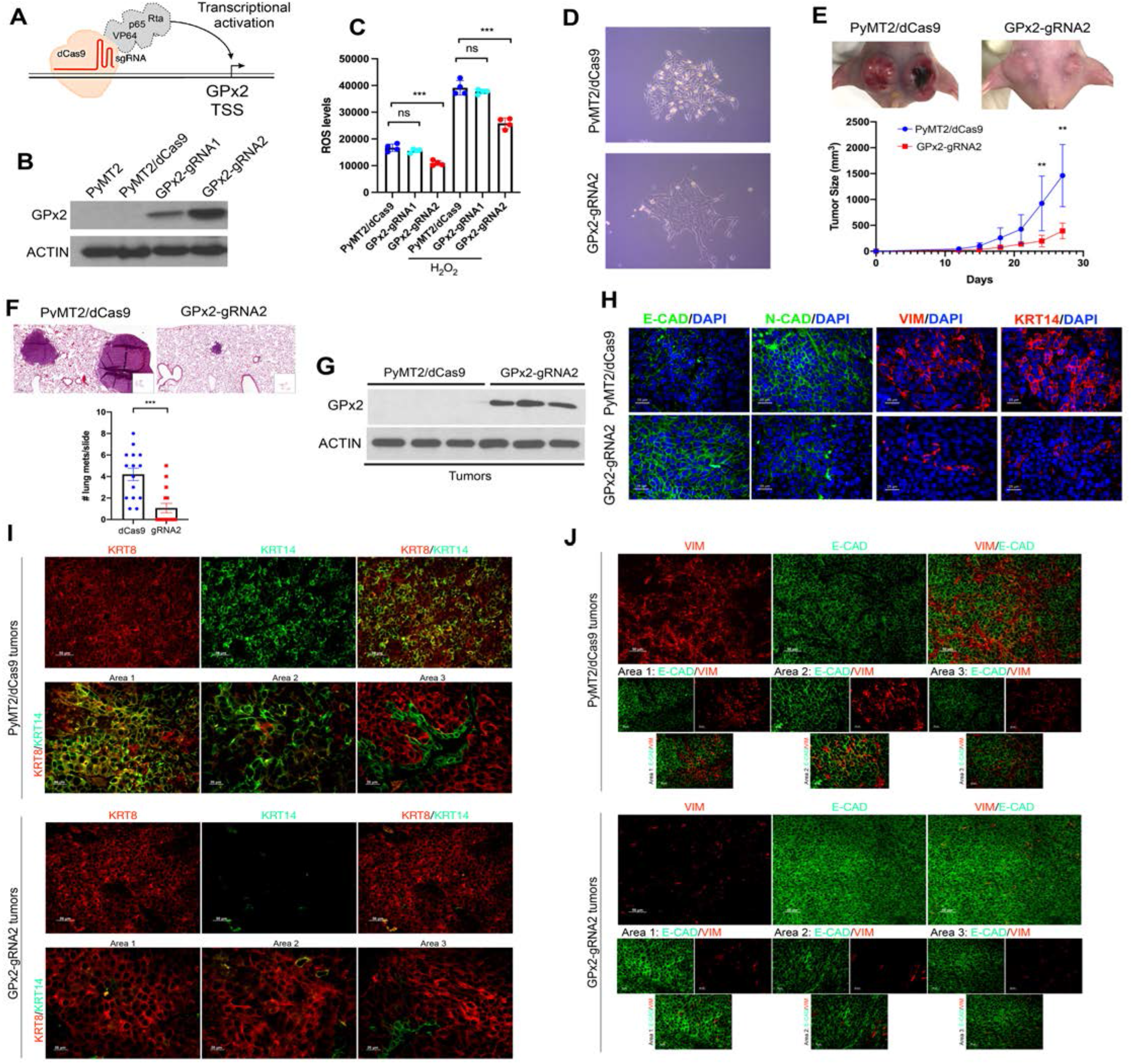
Endogenous induction of GPx2 expression in PyMT2 cells attenuates EMT dynamics. (**A**) Schematic representation of the sgRNA-dCas9-VPR complex. Transactivators VP64, p65, and Rta were directly fused with C-terminal of dCas9. sgRNAs bring dCas9 and transactivators to bind upstream of the GPx2 transcription start site (TSS) for activation of gene expression. (**B**) Two GPx2 targeting sgRNAs-dCas9-VPR were used to endogenously activate GPx2 expression in PyMT2 cells, and dCas9-VPR without sgRNA was used to control. Cell lysates were immunoblotted with anti GPx2 or Actin antibody. (**C**) ROS levels in PyMT2/dCas9 control and two clones of PyMT2/GPx2-gRNA are shown as Mean ±SEM; **p < 0.01, ***p < 0.001, ns indicates non-significance. (**D**) Phase-contrast images of PyMT2/dCas9 control cells and PyMT2/GPx2-gRNA2 cells at 10X magnification showing morphological difference between two cell lines. (**E**) PyMT2 control and GPx2-gRNA2 cells (1 × 10^6^) were bilaterally injected into mammary fat pads of female athymic nude mice (n=3 each group); representative images of tumor growth at 28 days post onset are shown (upper panels). Tumor growth curves over 28 days post tumor onset are shown as Mean ±SEM; **p < 0.01. (**F**) Scans of H&E stained sections of whole lung lobes (boxes) from mice carrying control or GPx2-gRNA2 tumors (upper panels). Graph showing number of metastatic foci in lung sections (n=5 sections per lung) from 3 mice per group; Mean ±SEM; ***p< 0.001. (**G**) Western blots show protein levels of GPx2 vs ACTIN in PyMT2/dCas9 control and GPx2-gRNA2 tumors (n=3 from independent mice). (**H**) Co-staining of E-CAD, N-CAD, KRT14, and VIM in PyMT2/dCas9 control and GPx2-gRNA2 tumors (n=6). (**I-J**) Immunofluorescent co-staining for KRT8 and KRT14 (I), or VIM and E-CAD (J) in PyMT2/dCas9 control vs PyMT2/GPx2-gRNA2 tumors (n=6) from 3 mice each.

### GPx2 overexpression in human aggressive breast cancer cells reverses the EMT dynamics

Lastly, to confirm these data in human breast cancer, we used a HER2-amplified cell line, JIMT1, which is GPx2 negative and more closely related to the luminal B subtype of the PyMT model. GPx2 OE in JIMT1 cells dramatically suppressed mammary tumor growth (**Fig. 5A-B**) and these effects were correlated with epithelial differentiation. Namely, compared to control JIMT1 tumors which were abundantly E-CAD^high^/VIM^high^ (**Fig. 5C**), GPx2 OE tumors were E-CAD^high^/VIM^low^ (**Fig. 5C**), thus shifting from an E/M to E state. These results demonstrate suppression of EMT by GPx2 in human BC cells, leading to the loss of E/M and M state, resulting in epithelial differentiation and malignancy suppression.

**Fig. 5.**
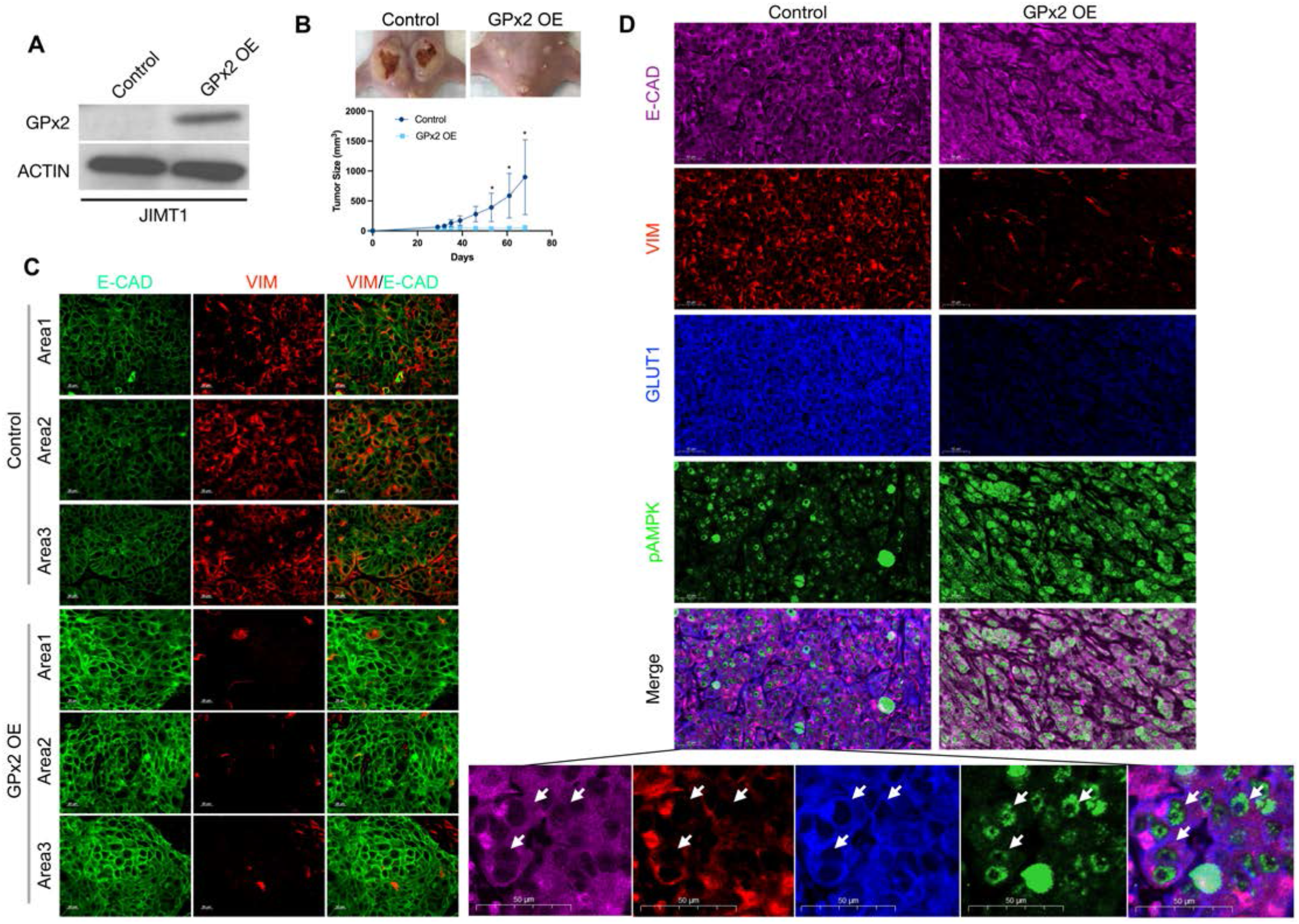
GPx2 overexpression in JIMT1 cells stimulates MET. (**A**) Immunoblotting analysis of GPx2 vs ACTIN in JIMT1 control and GPx2 OE cell lysates. (**B**) JIMT1 control and GPx2 OE cells (1 × 10^6^) were bilaterally injected into mammary fat pads of female athymic nude mice (n=3 each group); representative images of tumor growth at 68 days post onset are shown (upper panels); Mean ±SEM; *p < 0.05. (**C**) Immunofluorescent co-staining for VIM and E-CAD in GPx2 OE vs control JIMT1 tumors (n=6) from 3 mice each. (**D**) Co-staining for E-CAD/VIM/GLUT/pAMPK in GPx2 OE vs control JIMT1 tumors (n=6).

### HIF1α regulates metastasis via promotion of EMT dynamics

GPx2 knockdown in PyMT1 cells stimulates ROS/HIF1α signaling and metastasis (8), which might be due to activation of the EMT process. Others have shown that tumor cell intravasation was mediated by *Twist*, a *bona-fide* HIF1α target gene, which activates N-cadherin gene expression and EMT (32, 35). Alternatively, HIF1α transactivates the expression of *Snai1* or *Snai2*, which results in *Cdh1* gene repression (36-38). Interestingly, PyMT1/GPx2 KD tumors were shown to upregulate HIF1α, SNAI1, TWIST, and N-CAD while downregulating E-CAD protein expression (see **Fig. 3A-B**), which raises the question whether HIF1α promotes metastasis via EMT.

To test this hypothesis in the PyMT1/GPx2 KD model, we used echinomycin, a drug that inhibits the interaction of HIF1α with DNA and hence represses downstream gene transcription (39, 40). Treatment of PyMT1/GPx2 KD tumor bearing mice with echinomycin was shown to reduce mammary tumor growth by 50%, while partially renormalizing the tumor vasculature (8). To directly test effects on metastasis, mice were treated with echinomycin post-surgical excision of the primary tumor, to avert mouse morbidity by tumor burden as well as bypass the abnormal tumor vasculature that might impede drug perfusion (8). This also allows sufficient time for circulating tumor cells to build overt metastases in distant organs. One week post-surgery, mice were treated daily with intra-peritoneal injection of 10 μg/kg echinomycin for 7 consecutive days and then left untreated for 28 days to confirm effectiveness and durability of treatment. Remarkably, compared to vehicle-treated mice which produced a dramatic load of pulmonary metastases, echinomycin-treated mice were nearly devoid of metastases (**Fig. 6A-B**). Interestingly, echinomycin-treated tumors showed a dramatic reduction in SLUG, NOTCH1, and VIM expression relative to vehicle-treated tumors (**Fig. 6C**), and contained significantly less KRT8/KRT14 or E-CAD/VIM double-positive areas than vehicle treated tumors (**Fig. 6D-E**). These results underscore a predominant role for HIF1α in driving metastasis via EMT dynamics leading to the onset of E/M and M differentiation states.

**Fig. 6.**
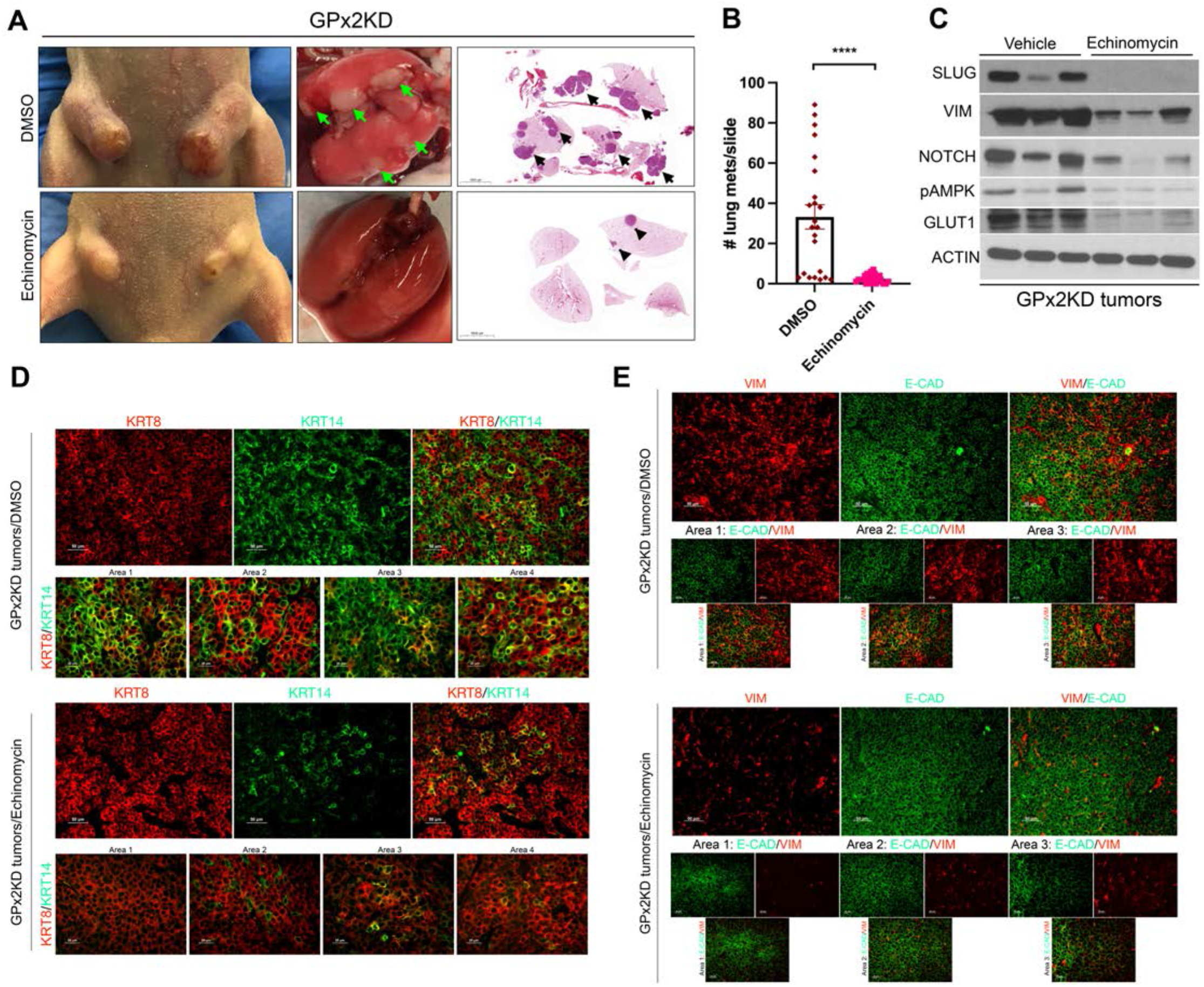
Inhibition of HIF1α suppresses spontaneous metastasis via changes in EMT dynamics. (**A**) Mice (n=10) bearing PyMT1/GPx2 KD tumors were treated post tumor onset with daily i.p injection of vehicle (DMSO) or 10 μg/kg of echinomycin/DMSO for 21 days after tumor growth to 64 mm^3^. Representative images of tumors from both groups are shown (left boxes). Mice then underwent survival surgery to remove primary tumor and were left untreated for one week to recover. Treatment was then resumed daily for one more week as above. One month later, mouse lungs were removed (second boxes), H&E-stained and scanned; scans of whole lung lobes (third boxes) from both groups are shown (3^rd^ boxes). (**B**) Quantification of lung foci from H&E stained sections are shown as Mean ± SEM; ****p < 0.0001. (**C**) GPx2 KD tumors from 3 independent mice that were treated with vehicle or echinomycin were subjected to immunoblotting with SLUG, VIM, NOTCH1, pAMPK, GLUT1, or ACTIN antibody. (**D-E**) Immunofluorescent co-staining of KRT8 and KRT14 (D), or VIM and E-CAD (E) in DMSO *vs* echinomycin treated PyMT1/GPx2 KD tumors from 3 mice each is shown.

### GPx2 knockdown results in phenotypic and metabolic adaptability

We previously showed that GPx2 KD promotes malignancy via metabolic modulation. Namely, GPx2 KD caused a shift from oxidative phosphorylation (OXPHOS) to aerobic glycolysis in most clusters (8). However, one of the clusters (cluster 5) was endowed with the ability to use both OXPHOS and glycolysis (8). Using phosphorylated-AMPK (pAMPK) and GLUT1 as surrogates for OXPHOS and glycolysis respectively, we were able to detect cluster 5-like cells in the GPx2 KD tumor by immunostaining (8). Interestingly, pAMPK and GLUT1 were both dramatically inhibited in echinomycin-treated GPx2 KD tumors (**Fig. 6C**), consistent with reports that HIF1α activates AMPK and glycolysis independently (41-44), implying HIF1α regulates metabolic plasticity.

To determine whether E, E/M, and M states spanning the EMT spectrum differed in their metabolic profiles, we co-stained tumors for epithelial markers (KRT8, KRT14; shown in yellow and red), and metabolic indicators (pAMPK, GLUT1; shown in green and blue). Remarkably, GPx2 KD tumors were enriched in carcinoma cells (mostly Area 1 but also Area 3) which were KRT8/KRT14/pAMPK/GLUT1 quadruple positive (**Fig. 7A;** right panels), relative to control tumors which were KRT8/pAMPK positive (**Fig. 7A;** left panels). Consistent with heterogeneity, Area 2 in the GPx2 KD tumor was represented by cells that were entirely KRT8/pAMPK positive (**Fig. 7A;** right panels). We conclude that E/M clusters adopt a hybrid metabolism, likely allowing them to switch between OXPHOS and glycolysis as they move from normoxic to hypoxic tumor niches. Alternatively, cells found at the boundary of normoxic (well-vascularized) and hypoxic (poorly-vascularized) tumor areas may use both metabolic modalities (42, 45-48).

**Fig. 7.**
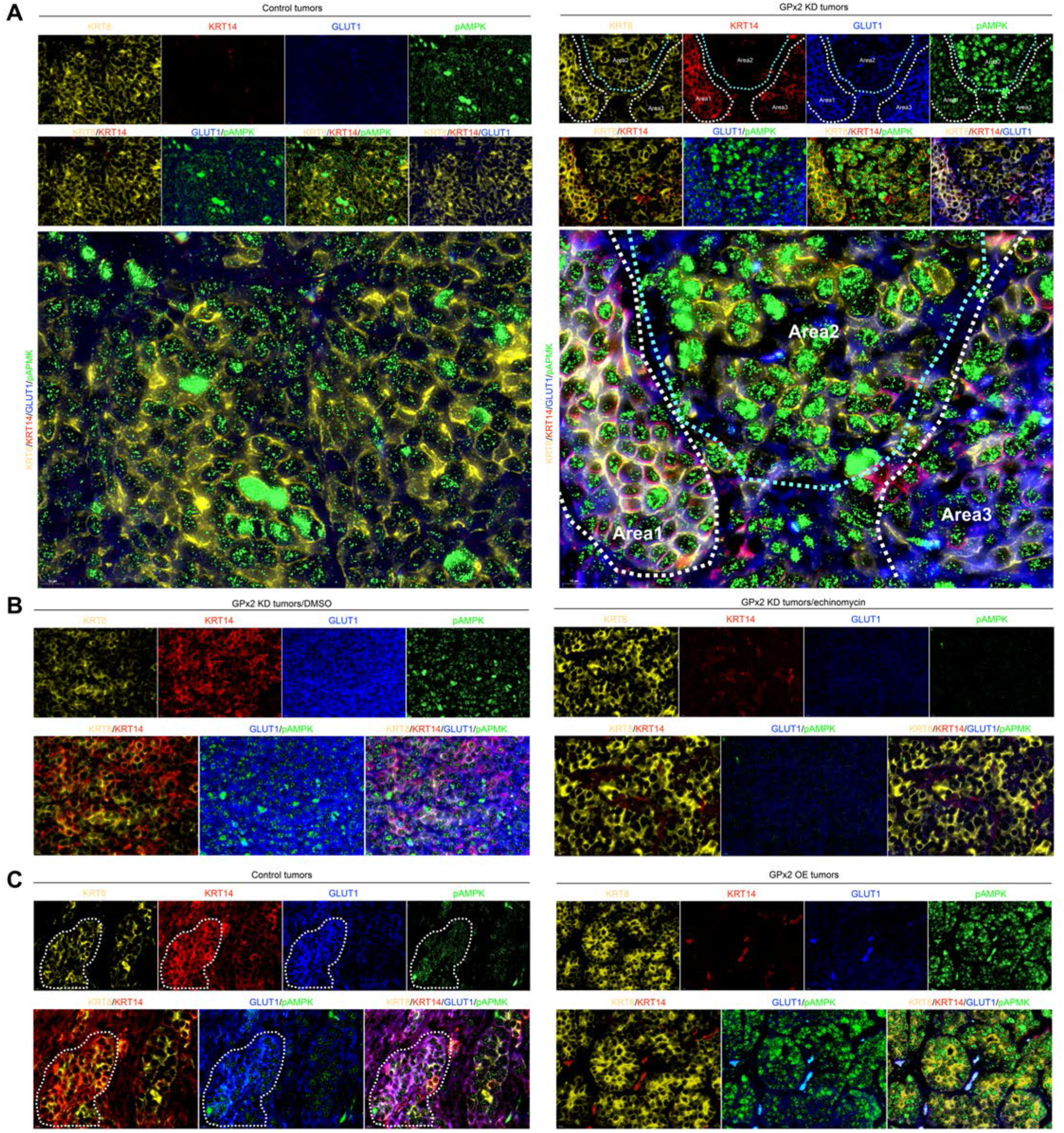
GPx2 KD results in E/M and OXPHOS/Glycolysis hybrid phenotypes that can be inhibited by GPx2 overexpression or HIF1α inhibition. (**A**) Co-staining for 4 markers consisting of KRT8 (yellow), KRT14 (red), GLUT1 (blue), and pAMPK (green) shows individual and overlay staining. Strong quadruple staining was observed in PyMT1/GPx2KD tumors especially in Area 1 and also in part of Area 3, but not in Area 2 which was KRT8+/pAMPK+ only (right panels). By contrast, control PyMT1 tumors were entirely KRT8/pAMPK positive (left panels). Staining was performed in 6 tumors from 3 mice each. (**B**) Co-staining for KRT8, KRT14, GLUT1, pAMPK in (B) PyMT2 control (left panels) and PyMT2/GPx2-OE (right panels) tumors; (n=6) from 3 mice; (**C**) Co-staining for the four above markers in GPx2 KD tumors (n=6) that were treated with DMSO (left panels) or echinomycin (right panels) from 3 mice.

Since HIF1α regulates pAMPK and GLUT1 expression, we tested whether echinomycin affects the expression of KRT8/KRT14/pAMPK/GLUT1 in tumors. Indeed, echinomycin-treated GPx2 KD tumors were markedly reduced in KRT14, GLUT1 and pAMPK expression, leaving tumors that were strongly KRT8+ (**Fig. 7B;** right panels) relative to vehicle-treated tumors which showed quadruple-stained areas (**Fig. 7B;** left panels). These data were in line with metastasis suppression by echinomycin, likely due to elimination of E/M and M cells in the GPx2 KD primary tumor or perhaps even in cells in circulation, consistent with enrichment of E/M cells in CTCs (49).

Next, we examined whether GPx2 OE in PyMT2 cells affects hybrid metabolism. Interestingly, PyMT2 tumors were strongly KRT8/KRT14/pAMPK/GLUT1 positive (**Fig. 7C;** left panels), supporting a role for E/M cells in metastasis. By contrast, GPx2 OE abolished KRT14/GLUT1 expression, resulting in strongly KRT8/pAMPK positive tumors (**Fig. 7C**; right panels). These data were consistent with malignancy suppression by GPx2 OE in PyMT2 cells (8). Similarly, compared to PyMT2/dCas tumors which were quadruple positive, endogenous upregulation of GPx2 suppressed KRT14/ GLUT1 positive cells, leaving only KRT8/pAMPK+ tumors (**Fig. S6A-B**).

Importantly, to confirm these data in human breast cancer cells, we stained the JIMT1 tumors for the four markers including E-CAD (Magenta), VIM (red), pAMPK (green) and GLUT1 (blue) (**Fig. 5D**). Indeed, a significant fraction of control JIMT-1 cells expressing E-CAD and VIM also expressed pAMPK and GLUT1 in the same cells (**Fig. 5D**, left panels; see arrows in bottom panels), indicative of hybrid metabolism. By contrast, GPx2 overexpressing JIMT-1 tumors showed dramatic reduction in VIM and GLUT1 expression, concomitant with increases in E-CAD and pAMPK, (**Fig. 5D**, right panels). These data were consistent with the notion that GPx2 suppresses mesenchymal and glycolytic genes, resulting in epithelial tumors overexpressing E-CAD and pAMPK, implying these tumors may use FAO/OXPHOS metabolism, consistent with other reports (50).

Collectively, these results strongly support the notion that GPx2 KD activates HIF1α signaling, enhancing mesenchymal gene expression, resulting in tumors enriched in phenotypic and metabolic hybrid phenotypes. Our data support the notion that the E-state utilizes OXPHOS; the M-state, glycolysis, and the E/M state uses both, implying that EMT and metabolism regulatory genes may influence each other expression to adapt to the higher metabolic demands of metastatic tumors (50).

## Discussion

Our findings highlight the far-reaching effects of GPx2 dysregulation on ROS/HIF1*α* signaling underlying EMT and metastasis. Our study provides novel insights into the diverse mammary tumor cell subpopulations (clusters) that are shaped by phenotypic and metabolic co-adaptation.

Single cell transcriptomics unraveled the dramatic effect of redox signaling by GPx2 KD on EMT, resulting in cells drifting through an EMT continuum, thus generating phenotypic and metabolic heterogeneity. Notably, GPx2 KD induced *de novo* expression of basal/mesenchymal genes (*Krt14, Twist2, Cdh2*) in all the luminal clusters while exacerbating the mesenchymal signature of cluster 3. This likely results in E/M clusters that may differ in their E to M gene expression ratio, thus creating a continuum of EMT states, with cluster 3 residing at the extreme mesenchymal transcriptional node. Interestingly, GPx2 KD increased the size of cluster 3 and decreased the size of luminal clusters 4/5, which could be due to gain of mesenchymal genes in the GPx2 KD tumor, converting cells in cluster 4 or 5 into cluster 3-like cells. This idea was corroborated by pseudotime trajectory analysis indicating closest proximity between cluster 5 and cluster 3, implying cluster 5 may reside in a highly hybrid E/M state within a continuum of states. In support of the E/M phenotype as a pivotal driver of metastasis, a recent study in the MMTV-PyMT model has shown that partial, but not full, EMT cells were enriched in lung metastases (16). Moreover, a study of pancreatic cancer heterogeneity identified early and late hybrid E/M clusters with the late clusters containing the most aggressive cells (17).

These studies argue against mesenchymal tumor cells as the drivers of metastasis, which implies that cluster 3-like cells may not actively promote metastasis (51). In fact, E/M tumor cells were found to be more efficient than M cells in promoting tumor initiating potential or metastasis, likely due to enhanced cell proliferation driving the self-renewal of E/M cells as compared to the more quiescent M cells (17). We thus speculate that cluster 3 cells reside at the apex of the CSC hierarchy, giving rise to progenitors or transit amplifying adopting an E/M state, hence supporting the positioning of cluster 3 at the root of the lineage trajectory.

Other studies have indicated that E/M carcinoma cells are not plastic but reside in a stable transcriptional hybrid state (15). We however speculate that E/M cells may convert into E or M cells dependent on contextual signals. It is conceivable that once E/M cells seed distant organs, they undergo MET into epithelial (E) cells in order to proliferate into metastatic colonies. Alternatively, E/M cells may convert into M cells in response to metastatic niche signals (e.g TGF*β* or HIF1*α*), thus favoring of a link between cancer stemness and tumor cell plasticity (52, 53).

In light of the dramatic inhibition of metastasis by echinomycin, we propose that HIF1α acts as a pivotal regulator of EMT dynamics. Namely, HIF1α may activate EMT via Snail upregulation, resulting in epithelial gene repression and E to M conversion. Otherwise, HIF1α may directly transactivate mesenchymal gene expression (not involving epithelial gene repression), thus generating an E/M hybrid state. For instance, HIF1α may transactivate *vimentin* or *Twist2*, which in turn activate N-cadherin (32, 54), thereby generating E/M hybrid clusters. It is tempting to speculate that echinomycin inhibits metastasis by eliminating E/M and M states, via suppression of M genes. Echinomycin reduced the level of E/M positive tumor areas coinciding with decreased KRT14 and VIM expression, resulting in luminal KRT8 or E-CAD enriched tumors. Indeed, echinomycin was used to target hematopoietic stem cells (40), suggesting that HIF1α-based therapy may inhibit stem-rich or metastasis-prone tumors.

Other than promoting the E/M state, GPx2KD/HIF1*α* stimulates tumor metabolism, and these two processes appear to be tightly interwoven (47, 48). While most tumor clusters used aerobic glycolysis (Warburg effect), cluster 5 was shown by IPA to have the potential to use both OXPHOS and glycolysis (8). In support of HIF1*α* as a central regulator of EMT and metabolism, echinomycin was able of inhibiting the expression of mesenchymal VIM, SNAIL, and NOTCH1 proteins as well as pAMPK and GLUT1 metabolic markers that are indicative of OXPHOS and glycolysis (see **Fig. 6C**) (47, 48). By contrast, treatment of GPx2 KD tumors with lapatinib, an inhibitor of ErbB1/2, which was hyper-phosphorylated in GPx2 KD tumors, had no effect on metastasis (unpublished data). Other than targeting E/M cells in the primary tumor, echinomycin may obliterate CTCs which contain E/M cells that also use OXPHOS and glycolysis, thus explaining the dramatic effect of metastasis inhibition by echinomycin (42, 49, 55).

We further predict that HIF1*α* regulates the interplay between E/M and hybrid metabolic states. Namely, HIF1*α* stimulates the expression of glycolytic enzymes as well as AMPK (41-44). AMPK was shown to promote fatty acid uptake via manipulation of fatty acid translocase (FAT/CD36), resulting in fatty acid oxidation (FAO) and hence OXPHOS (56-58). We speculate that by promoting EMT dynamics, glycolysis, and possibly FAO via AMPK, HIF1*α* fosters a mutual relationship between E/M state and hybrid metabolism. Interestingly, TGF*β*, a strong EMT inducer, was shown to activate glycolysis as well as FAO, which in turn produces acetyl-CoA that feeds into OXPHOS, an effect which was mimicked by SNAI1 overexpression (59). In further support of our data, others have shown that breast M and E (more likely E/M) CSCs which were low-ROS and high-ROS accordingly, utilize OXPHOS and glycolysis/OXPHOS respectively (22, 60). These findings are in agreement with GPx2 KD, which mimics a high-ROS state that may contribute to the hybrid state, perhaps by activating HIF*α* and AMPK and thus glycolysis and FAO (OXPHOS), thereby consolidating a phenotypic and metabolic reprogramming, underlying metastasis. However, the exact mechanism for maintaining a balance between glycolysis and FAO in the hybrid state warrants further investigation.

Altogether, our findings provide a conceptual framework to unravel mechanisms and/or molecular players that regulate the phenotypic and metabolic co-dependence in tumor cells with the potential to yield novel therapeutic strategies in metastatic breast cancer.

## Material and Methods

### Cell lines

Primary mammary tumor cell lines were generated from PyMT transgenic mouse tumor model as following procedures. Tumors were excised, washed in PBS and diced with a razor blade into smaller fragments. Each gram of tumor was incubated with 10 mL of digestion media (1 mg/mL Collagenase type I, 100 unit/ml Hyaluronidase, 100 units/ml Penicillin/ streptomycin, 2 mg/ml of BSA in Medium 199). The suspension was incubated at 37 °C for 3 hrs with occasional mixing. Digested material was spun at 1000 rpm for 5 min and pelleted cells were plated overnight at high density (1×10^7^ cells per 10 cm dish) in DMEM/20%FBS supplemented with 10 μg/ml insulin and 20 μg/ml EGF. Fibroblasts were depleted from epithelial tumor cells by limiting trypsinization. Epithelial cells were grown in culture for several weeks till they reached crisis and adapted to in vitro culture conditions. The human MDA-MB-231 and JIMT1 breast cancer cell lines were cultured in DMEM medium (Gibco) supplemented with 10% FBS and 1% Pen-Strep (Gibco).

### Animal studies

Mice were housed and maintained by the Animal Studies Institute at the Albert Einstein College of Medicine. Animal protocols used for this study were reviewed and approved by the Institute for Animal Studies. The mice were obtained from Jackson laboratories. MMTV-PyMT transgenic mice that were used in another study (61) were derived by brother-sister (sibling) mating of progeny from a cross of p21 knockout female (Bl6/129S/background) with MMTV-PyMT heterozygous male (FVB background). Only female p21wt/wt PyMT mice derived from F2 progeny of selected brother sister matings were used to derive primary mammary tumor cell lines. These resulting recombinant mice were on a mixed Bl6/129S/FVB/N background that were backcrossed in the FVB/N background. PyMT xenograft models were generated by injection of p21wt/wt PyMT cell lines derived from the same mammary tumor (PyMT1 and PyMT2) into female athymic nude mice obtained from Jackson laboratories. For MDA-MB-231 xenografts, 1 million cells suspended in 200 ul PBS containing 25% Matrigel were bilaterally injected in the flanks of athymic female nude mice. For JIMT1 xenografts, 1 million cells suspended in 200 ul PBS were bilaterally injected into the flanks of athymic female nude mice.

### Tumor growth kinetics

Tumor growth curve was determined by measuring tumor size. The tumor size was measured twice a week using a caliper. Tumor volume was determined upon the formula: tumor volume = shorter diameter^2^ x longer diameter/2. Mice (three per group for PyMT2/dCas vs. PyMT2/dCas-GPx2-gRNA2; ten for PyMT1/GPx2 KD treated with vehicle vs. ten for PyMT1/GPx2 KD treated with echinomycin; three per group for JIMT1 control vs. JIMT1/GPx2 OE) were sacrificed at end point. Data are displayed as mean tumor volume -/+ SEM. Statistical analyses were performed comparing individual time points by unpaired t-test and significant differences were established as p-value < 0.05.

### Lung metastasis

Mice bearing mammary tumors were sacrificed at end point of 1-2 cm diameter, as allowed by our animal protocol. Lungs were inflated by tracheal cannulation with injection of 1-2 ml of 10% neutral buffered formalin. Formalin fixed lungs were paraffin embedded and blocks sectioned on a tissue microtome (Leica Microsystems) at 5 μm. Lungs were serial sectioned through the tissue and sets of 5 serial sections, at 300 um intervals. Analysis was performed on whole sections after it was determined by inspection that metastases seeded in random locations. Data are displayed as mean foci number ± SEM. Statistical analyses were performed using unpaired t-test and significance determined at p < 0.05.

### Constructs

The third generation lentiviral transfer plasmid pXPR_dCas9-VPR_sgRNA was cloned from pXPR_dCas9-VP64-Blast (Add-gene plasmid #61425) (62). For mouse GPx2 sgRNA design, two custom anti-sense DNA oligonucleotides (CTTTGTTCAGTGGCAGTAAG, TTGTTCAAACAGTTCACAGG) were annealed and ligated into pXPR_dCas9-VPR_sgRNA. Non-targeting pXPR_dCas9-VPR with no GPx2 sgRNA inserted recognition sequence was used as a control. Mouse GPx2 plasmid (GenScript) was subcloned by PCR into Xho1/BamH1 restriction sites of lentiviral expression vector pLVX-puro (Clontech), using 5’ region primer: 5’-TAT CTC GAG GCC ACC ATG GCT TAC ATT GCC AAG TCG-3’ and 3’ region primer: 5’-TAT GGA TCC CTA GAT GGC AAC TTT GAG GAG CCG-3’. Lentiviral particles produced from the obtained mGPx2-pLVX-puro plasmid were used for expression of mGPx2 protein in the PyMT2 mammary tumor cell line.

### Lentivirus production

Lentiviral particles were generated by transient co-transfection of 293T cells with lentivirus-based vector expressing either the full-length clone of desired gene or an shRNA sequence to target-specific RNA of gene. Briefly, a 100-mm dish seeded with 3×10^6^ cells were transfected with 0.6 μg of lentiviral packaging gene TAT, 102 RVE, and GAG/POL and 1.2 μg of VSV-G and 12 μg of DNA of interest in lentiviral backbone. Fugene was used to transfect the cells. Following 48 hours of transfection, supernatant was collected, centrifuged at 2,000 rpm for 10 minutes, and filter sterilized.

### GPx2 lentivirus overexpression

The open reading frame of mouse GPx2 (GenScript NM_030677.2) was subcloned by PCR into Xho1/BamH1 restriction sites of lentiviral expression vector pLVX-puro (Clontech), using 5’ region primer: 5’-TAT CTC GAG GCC ACC ATG GCT TAC ATT GCC AAG TCG-3’ and 3’ region primer: 5’-TAT GGA TCC CTA GAT GGC AAC TTT GAG GAG CCG-3’, and the open reading frame of human GPx2 (GenScript NM_002083.4) was subcloned by PCR into Xho1/BamH1 restriction sites of lentiviral expression vector pLVX-puro (Clontech), using 5’ region primer: 5’-CCC CTC GAG ATG GCT TTC ATT GCC AAG TCC TTC TAT GAC-3’ and 3’ region primer: 5’-CCC GGA TCC CTA TAT GGC AAC TTT AAG GAG GCG CTT GAT-3’. Recombinant lentiviruses expressing mGPx2 and hGPx2 were packaged in 293T cells and expressed in cells as described above.

### Lentiviral transduction

Cells were seeded at 1×10^5^ per 12-well plate for 24 hrs, treated with 250 μl viral solution containing 10 μg/ml polybrene for 1 hour and incubated in DMEM/10% FBS without antibiotics for 24 hours. Cells were expanded into a 10-cm dish with selective antibiotics.

### Antibodies

Antibodies against GPx2, HIF1α, NOTCH, pAMPK(staining) were obtained from Abcam. Mouse monoclonal anti KRT14(WB), KRT18, and rabbit polyclonal anti TWIST, EphA2 were obtained from Santa Cruz Biotechnology. Rat anti KRT8(staining) was purchased from DSHB and Rat anti E-cad (staining) was from Santa-Cruz Biotechnology. Rabbit anti VIM, SLUG, E-CAD, pAMPK(WB), and mouse anti SNAI1 were from Cell signaling and β-actin antibody was from Sigma.. Rabbit anti KRT14 antibody (staining) was from Biolegend. Guinea pig anti KRT14 or anti VIM antibody (staining) was from Progen. Mouse anti N-Cad was from BD Transduction Labs. Rabbit anti-SLC2A1(GLUT1) antibody was purchased from BioSource.

### Immunoblotting

Cells or tissues were extracted in RIPA solubilization buffer (50 mM Tris-HCL pH 7.5), 150 mM NaCl, 0.5mM MgCl2, 0.2mM EGTA, 1% Triton X-100) including protease and phosphatase inhibitors. 30 μg protein were loaded on 7-12% SDS-polyacrylamide gels and transferred to Immobilon membranes. Blots were probed overnight at 4 °C with indicated antibodies and developed by chemiluminescence (Perkin Elmer).

### Immunofluorescence

Formalin-fixed/paraffin-embedded tumor tissues were sectioned in 5 μm thickness, deparaffinized in xylene, and rehydrated in a series of 100% ethanol, 95% ethanol, and distilled water. Antigen retrieval was performed 1X antigen retriever solution (Sigma, pH 6.0). Tissues were incubated with primary antibody in 5% donkey serum, 2% BSA, 0.5%TX-100 in TBS, followed by incubation of Alexa Fluor secondary antibody for 1.5 hr at room temperature, washed and counterstained with DAPI to visualize nuclei.

### ROS Measurements

ROS detection reagent, 2′,7′-dichlorofluorescein (DCF) (Invitrogen, C6827) was used to determine intracellular ROS levels. DCF was reconstituted using anhydrous dimethylsulfoxide (DMSO, Sigma) with optimal working concentration at 10 μM in DMEM/2% serum. Media with DCF were added to cells in 96 wells and incubated for 1 hr in the dark at a 37 °C, 5% CO_2_ incubator. After incubation, media with DCF was removed and cells were washed twice with PBS. DCF fluorescence intensity relative to background fluorescence in wells that do not contain cells was determined using a fluorescent microplate reader using excitation wavelength at 495nm and emission wavelength at 520nm. The results were normalized to cells/well. Data are displayed as Mean H2DCFDA fluorescence intensity ± SEM.

### TOP/FOP flash assay

Cells were plated in 24-well plates in duplicates. Reporter plasmid TOP-Flash or FOP-Flash, which contain three optimal copies of the TCF/LEF-binding site (TOPFlash), or mutated copies of the TCF/LEF-binding site (FOPFlash) upstream of a minimal thymidine kinase promoter directing transcription of a luciferase gene were used. (TOPFlash) or (FOPFlash; 0.5 mg) together with a Renilla luciferase plasmid (0.1 **m**g) were transfected using Lipofectamine LTX (Invitrogen) according to the manufacturer’s protocol. Cell lysates were obtained using lysis buffer provided in Dual Luciferase Assay Kit (Promega). Firefly and Renilla luciferase readings were recorded 48 hours posttransfection using Luminometer (Promega). The firefly luciferase activity is then normalized to the Renilla luciferase activity, and fold increase in TOP-Flash activity compared with FOP-Flash was plotted as mean -/+ SEM of triplicate tests and validated by t-test, P **<** 0.05.

### Cell isolation of mammary tumor cells for single cell RNA sequencing

PyMT1 control and PyMT1/GPx2 KD tumor (one tumor each) generated by injection of GFP-labelled tumor cell lines, was excised and mechanically chopped with scalpels, incubated in DMEM/F12 with 5% FBS, 5 μg/ml insulin, 2 mg/ml collagenase, and 150 μg/mL DNase at 37 °C for 45 min. Cell clamps were washed and dissociated with 2 ml of 0.05% trypsin/EDTA at 37 ^°^C for 10 min and trypsinization stopped with 10ml PBS-2%FBS-EDTA. The samples were filtered through a 70 μm cell strainer, pelleted at 1200 rpm at 4 ^°^C for 5 min, and resuspended in 1 ml of 1x red blood cell (RBC) lysis buffer and 10ml PBS-2%FBS-EDTA was added to stop lysis. Cells pellets were resuspended in 1ml PBS-2%FBS-EDTA and viability-controlled by trypan blue dye exclusion.

### Flow cytometry

GFP-labelled cells from PyMT1 control and PyMT1/GPx2 KD tumor were sorted using a FACSAria II (BD Biosciences). Singlets were selected by using standard forward scatter width versus area. Cell viability was controlled by DAPI staining and single cell suspensions were sorted in DMEM/10% FBS.

### Library preparation and single cell RNA sequencing

Library preparation was performed by Mr. David Reynolds at the AECOM genomic core facility. GFP-sorted single cell suspensions from one control PyMT1 tumor and one PyMT1/GPx2 KD tumor containing at least 80% viable cells were processed to capture at least ∼10,000 viable cells per sample. The libraries were prepared (individual lanes on the 10X Chromium) using the 10X Single Cell 3’ v3 kit using ∼10,000 cells per lane on the 10X Chromium microfluidics device (10X Genomics, Pleasanton, CA). The qualified libraries were sequenced on Illumina Hiseq 6000 platform (Novogene Corp, Sacramento, CA), with a standard paired-end 150bp (PE150) at each end.

### Processing of scRNAseq data

The original data obtained from Illumina® HiSeq platform (Novogene Corp, Sacramento, CA), were transformed to sequenced reads by base calling. Raw data obtained as FASTQ files which contain sequenced reads were subjected to quality control using OpenGene’s fastp v0.19.4 preprocessor (63). Cellranger 3.0.2. software was used to provide reads alignment and gene annotation to the mm10 mouse reference genome. The STAR aligner employed within cellranger 3.0.2 was used to perform alignment. Fragments Per Kilobase of transcript per Million mapped reads (FPKM) values were quantified using each cell-barcode in combination with unique molecular identifiers (UMIs) using the default cellranger. The quantified UMI counts were used to form an unfiltered gene-barcode matrix and exported for further downstream analysis.

### Quality control and normalization

The unfiltered gene-barcode matrix for each sample was imported into R 3.6.1 and converted into SingleCellExperiment (SCE) objects. Any barcodes with a total UMI count greater than 200 were classified as cells. Seurat 3.1.1 was used for further quality control and preprocessing of the exported SCE objects. Cells with a minimum cut-off of 200 and a maximum cut-off of 4500 total RNA count were filtered. In addition, cells with a percentage of total reads that aligned to mitochondrial genome greater than 10% were removed (inferred as cells undergoing stress and cell death). After quality control, a total of 7621 cells from PyMT1 control tumor and 7768 cells from PyMT1/GPx2 KD tumor were used for normalization and downstream analyses. Cells passing QC were normalized using the SCTransform pipeline in Seurat 3.1.1.

### Integration and clustering

After normalization, the data obtained from PyMT1 control and PyMT1/GPx2 KD tumor underwent comprehensive integration using the integration pipeline in Seurat with the genes commonly retained between control and GPx2 KD tumor serving as integration anchors (64). Principal component analysis was performed to enable unsupervised clustering on the integrated data from PyMT1 control and PyMT1/GPx2 KD tumor, and clustered cells were projected in two dimension using Uniform Manifold Approximation and Projection (UMAP) for cluster visualization (65) Louvain community detection algorithm was further applied to identify the cluster partitions, with a resolution parameter of 0.4 that was identified by building a clustering tree on the datasets to optimize the number of clusters.

### Cluster identification and annotation

For cluster identification in the integrated dataset, we performed manually supervised-automated methods of cell-type identification. Canonical cell-type marker genes for basal (e.g, Krt5, Krt14) and luminal cells (e.g. Krt8, Krt18, Cldn) were annotated to identify which clusters belonged to which cell type. Seven epithelial clusters (cluster 0, 1, 2, 3, 4, 5, 6), and two non-epithelial clusters (cluster 7 and 8) were identified. Clusters 7 and 8 were tumor associated stromal cells that might be co-sorted with GFP+ tumor cells.

### Cluster comparison and visualization

To compare cluster specific genes under two conditions, we used feature plot and violin plot functions to highlight expression patterns of marker genes of interest. Dot plots were used to visualize the average expression pattern of percentage and intensity of expression of genes of interest cells across all cell clusters.

### Cell trajectory analysis

Pseudotime cell trajectory analysis was performed using Monocle3 package (https://cole-trapnell-lab.github.io/monocle3). The batch corrected and integrated data between control and GPx2 KD tumors using Seurat were used as input to Monocle3. The top 2000 highly variable genes in the integrated dataset were used to compute the pseudotime ordering and the cell trajectory analysis. Cells were plotted in UMAP space for visualization of pseudotime trajectories after clustering. We manually placed cluster 3 (basal/mesenchymal-like cell population) as root population by specifying the root_cells parameter. We further examined some basal/luminal/epithelial/mesenchymal genes found to be differentially expressed across the pseudotime trajectory.

### Differential gene expression analysis

To investigate gene expression change by cluster comparison between PyMT1 control and PyMT1/GPx2 KD tumor, a joint variable of condition * cell-type was used to calculate cluster-type specific differential expression. Genes with an adjusted p-value less than 0.05 were classified as significantly differentially expressed. The differential gene expression analysis was carried out using a Wilcoxon Rank sums test with Benjamini-Hochberg test for multiple testing correction, with the default parameters of the *FindMarkers()* function in Seurat.

### Ingenuity Pathway Analysis

To investigate overrepresented pathways in cluster 3, the differentially expressed genes between cluster 3 and the rest clusters 0, 1, 2, 4, 5, 6 were uploaded to Ingenuity Pathways Analysis (IPA) www.ingenuity.com), for core analysis, and a p-value of 0.05 was used as a cut-off to determine statistically significant enrichment of a pathway or annotated gene grouping present in the Ingenuity Knowledge base. A Z-score greater than +2 or less than -2 was used as a cut-off to predict a pathway activation or inhibition, respectively.

### Statistical Analysis

Results represent the Mean ± SEM (standard error of the Mean) for indicated experiments with at least 3 independent biological replicas. The statistical methods used are described in the figure legends. Most data were analyzed by the unpaired two-tailed Student’s t test to compare two groups with significance set at the p value < 0.05. Statistical analysis was performed using the Prism 9 (GraphPad) software.

## Acknowledgments

This work was supported by grants from Breast Cancer Research Foundation (BCRF), and Cure Breast Cancer Foundation (CBCF) to (R.B Hazan and L. Norton). BCRF was funded by generous gifts from Jane and Neil Golub. Zuen Ren was in part supported by a predoctoral fellowship grant from China. We are grateful to Mr. David Reynolds at the genomic core facility, the histology core and the NCI Albert Einstein Cancer Center grant (P30CA013330). We are greatly thankful to Dr. Ralph Garippa, Director of the Gene Editing and Screening (GES) Core Facility at Memorial Sloan Kettering Cancer Center, NY and to the Core Facility team for their services.

## Author contributions

R.B.H and ZR conceived the project, designed the study and wrote the original manuscript. Z.R. executed a majority of the experiments under the supervision from R.B.H., Z.R. and H.L. performed in vivo studies and analyzed the data. K.S. made and expressed the mouse GPx2 construct into PyMT2 cells. Z.R. made and expressed the human GPx2 construct into JIMT1 cells. O.B. expressed CRISPR/dCas 9 system construct into PyMT2 cells and analyzed ROS effects. Z.R. performed the single cell RNA-seq with help from H.L., K.S., O.B. for sample prep. Z.R., M.D., A.T.F. and J.C.M. carried out the computational analysis of the single cell RNA sequencing data. Z.R., M. M.G and O.B. performed Western Blots of cells and tumor lysates and analyzed changes in EMT. M.P. contributed to the conceptualization of scRNA-seq. L.N was involved in the conceptualization and discussion of the results. M.D., J.C.M., M.P., LN were involved in editing the manuscript.

## Data and Materials Availability

The single cell RNA sequencing data reported in this manuscript are available at https://www.ncbi.nlm.nih.gov/geo with access number GSE152368. Coding analyses for single cell RNA sequencing data are available in GitHub with the URL accession link https://github.com/Malindrie/Breast-cancer-scRNA-seqanalysis.

