## Supplemental Figures for "Redox signaling regulates breast cancer metastasis via HIF1α-stimulated EMT dynamics and metabolic reprogramming"

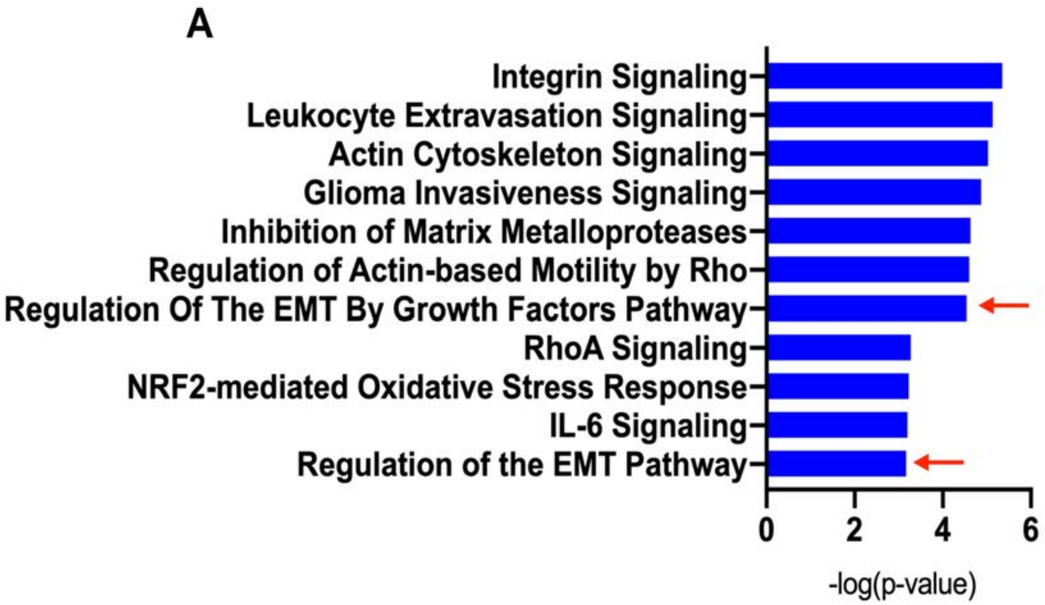


**Fig. S1. Ingenuity Pathway Analysis confirms that cluster 3 is basal/mesenchymal-like population overrepresented with EMT and stem like pathways.**

(**A**) Core analysis of differentially expressed genes between cluster 3 and clusters 0, 1, 2, 4, 5, 6, revealed overrepresented pathways that were enriched in cluster 3.

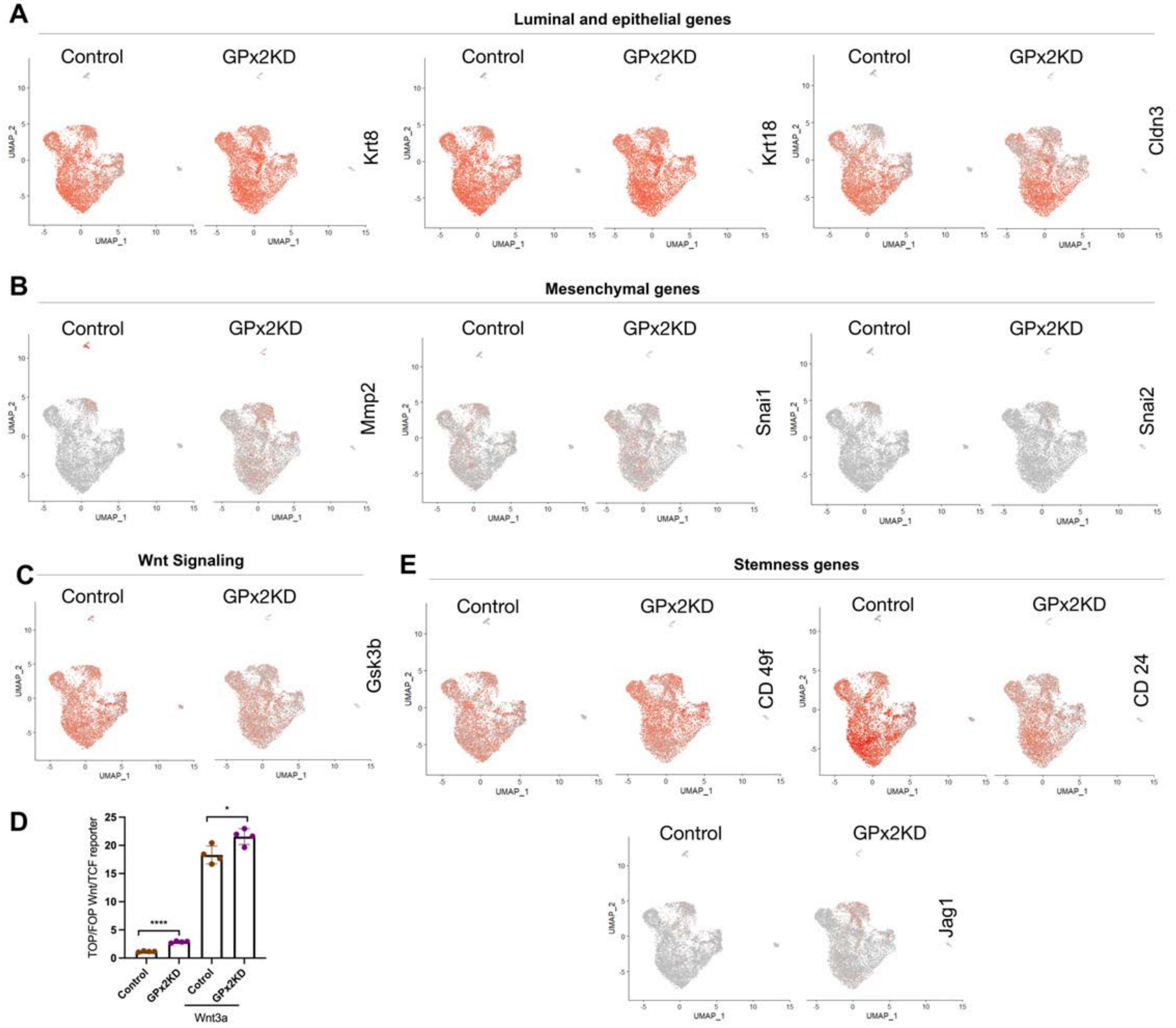


**Fig. S2. Feature plots highlight expression changes of EMT- and stemness-associated genes in PyMT1/GPx2 KD relative to control tumor.** (**A**) Feature plots in low-dimensional space showing expression of luminal and epithelial marker genes *Krt8*, *Krt18*, *Cldn3* in the GPx2 KD relative to control tumor. (**B**) Feature plots showing expressions of mesenchymal genes, *Mmp2*, *Snai1*, and *Snai2* comparing GPx2 KD to control tumor. (**C**) Feature plots showing expression of *Gsk3*β in the GPx2 KD relative to control tumor. (**D**) TOP/FOPFlash assay of GPx2 KD vs control cells that were untreated or treated with Wnt3a condition media for 24 hr under hypoxia (1% $O_2$). (**E**) Feature plots showing expression of stemness marker genes *CD49f*, *CD24*, and *Jag1* in the GPx2 KD relative to control tumor.

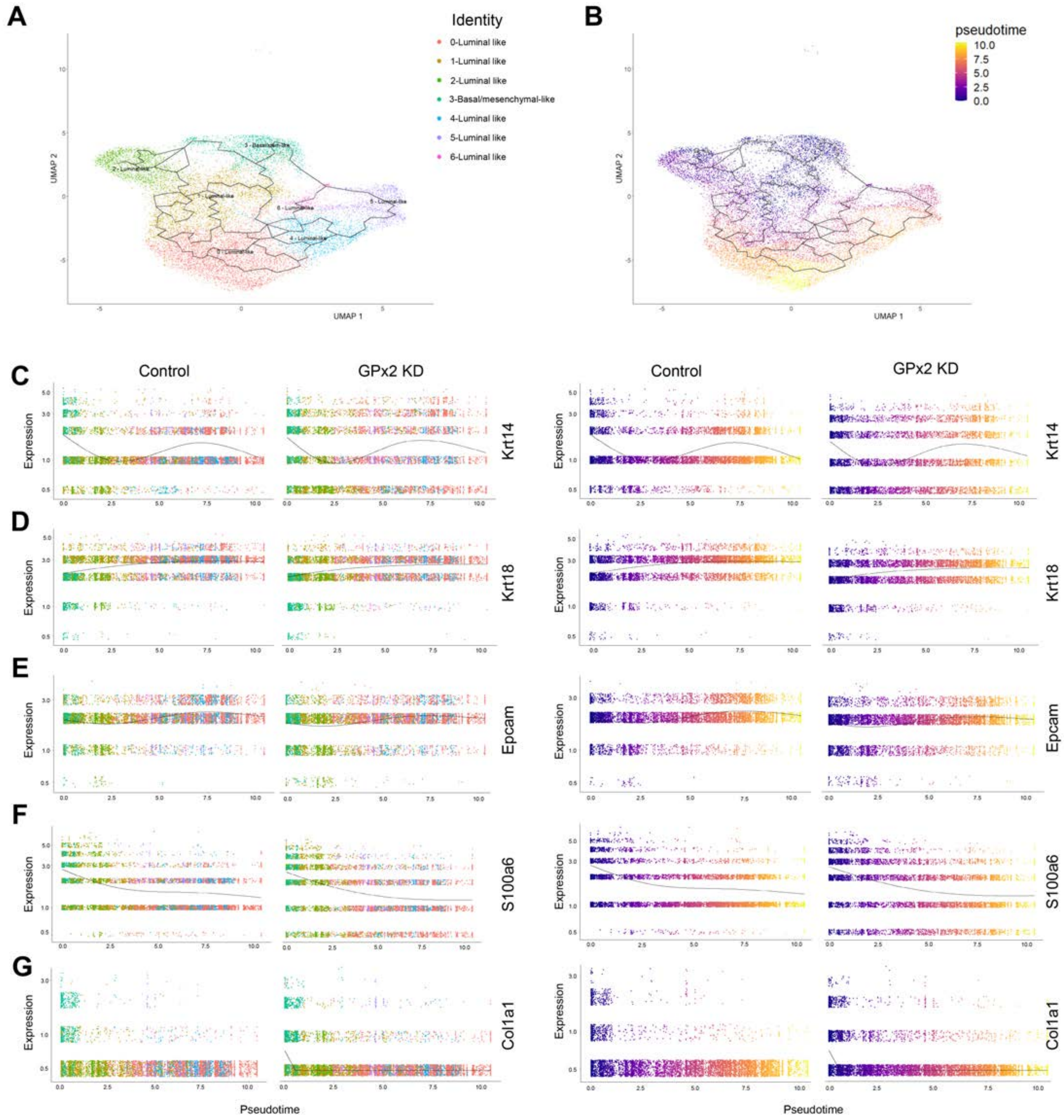


**Fig. S3. Cell trajectory analysis highlights a transcriptional EMT continuum across the tumor clusters.** (**A**) UMAP projection of six luminal-like clusters, one basal/mesenchymal-like cluster from comprehensively integrated clustering results, shared by one PyMT1 control and one PyMT1/GPx2 KD tumor. (**B**) Pseudotime time trajectories projected onto UMAP in (A); Color reflects pseudotime distance (basal/mesenchymal-like population located at t = 0). (**C-G**) Gene expression levels of indicated genes (*Krt4/18*, *Epcam*, *S100a6*, *Col1a1*) in each individual cell over pseudotime in control and GPx2 KD tumor; color displays each individual cell corresponding to individual clusters (left two columns), and color reflects pseudotime distance (basal/mesenchymal-like population located at t = 0) (right two columns).

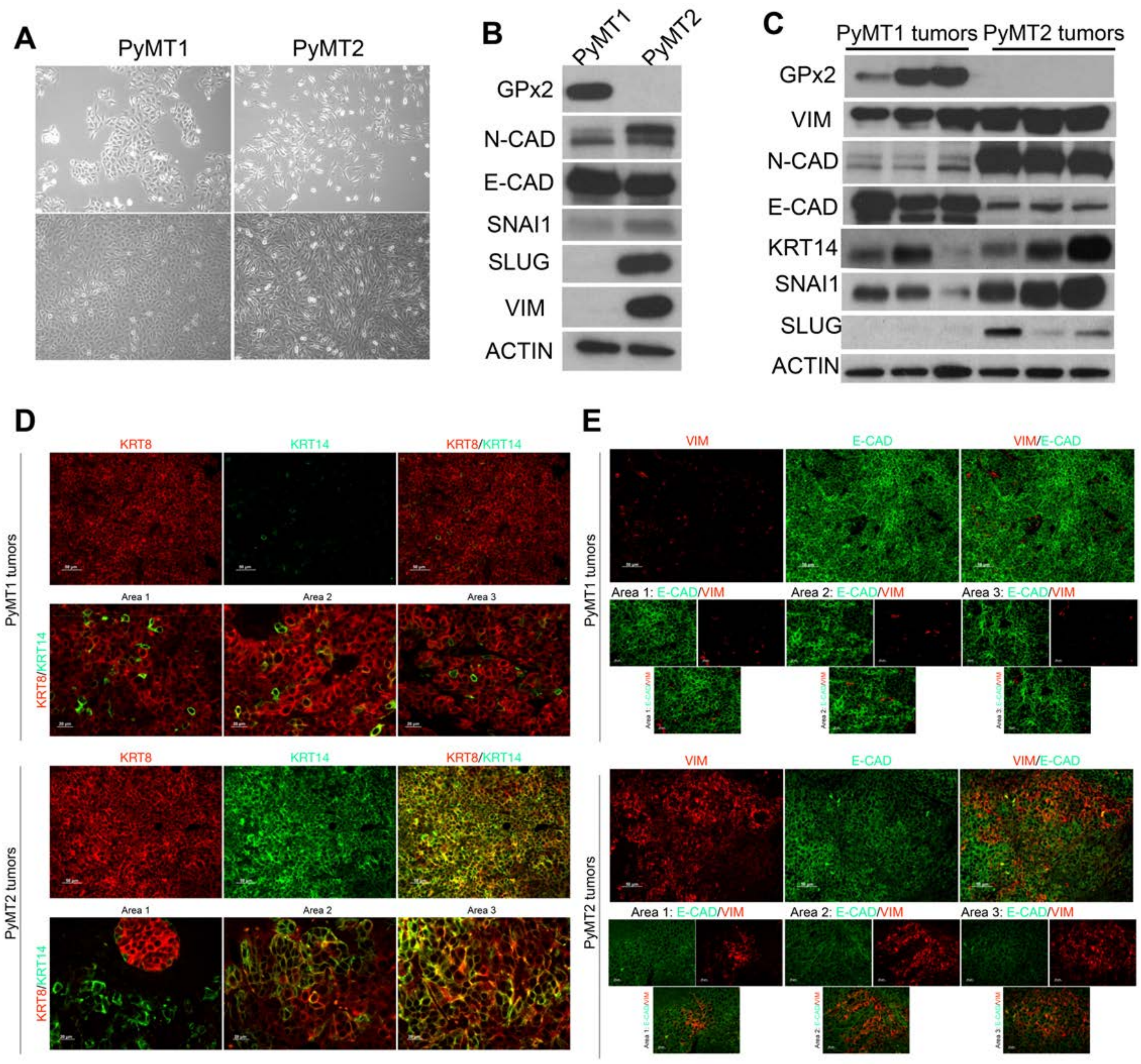

**Fig. S4. Correlation between metastatic potential of PyMT cell lines and EMT dynamics.**

(**A**) Phase-contrast images of non-metastatic PyMT1 cells and metastatic PyMT2 cells (10X magnification) at high and low dilution, show an epitheloid vs a spindle-like morphology respectively. (**B**) Western blots showing expression of GPx2, N-CAD, E-CAD, SNAI1, SLUG, VIM vs ACTIN in PyMT1 cells and PyMT2 cells. (**C**) Western blots of PyMT1 and PyMT2 mammary tumor lysates showing expression of GPx2, VIM, N-CAD, E-CAD, KRT14, SNAI1, SLUG, vs ACTIN (n=3; 3 independent mice). (**D-E**) Immunofluorescent co-staining analysis of KRT8 and KRT14 (D) or VIM and E-CAD (E) in PyMT2 vs PyMT1 tumors (n=6) from 3 mice each.

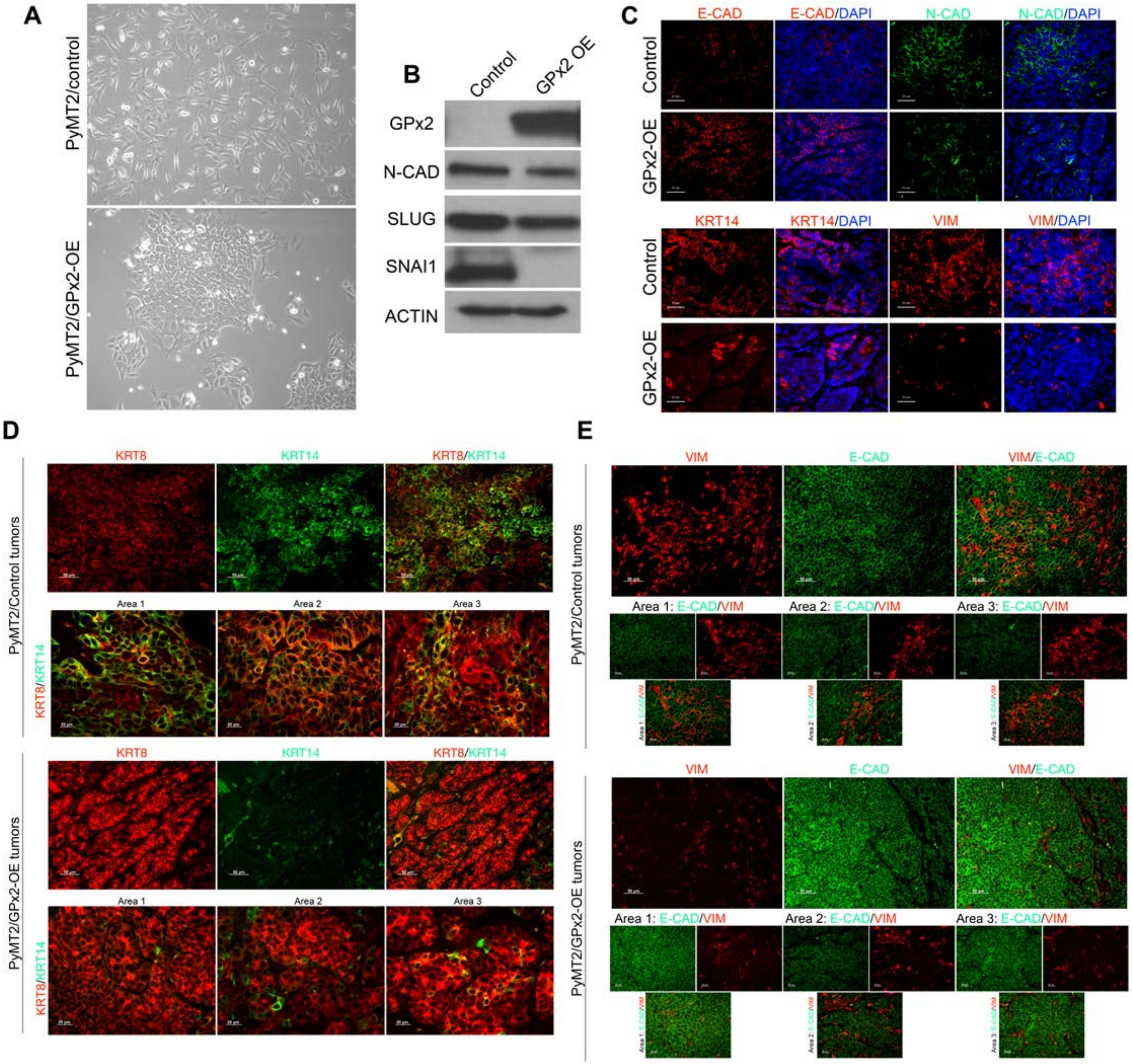

A
PyMT2/control
PyMT2/GPx2-OE
B
Control
GPx2 OE
GPx2
N-CAD
SLUG
SNAI1
ACTIN
C
E-CAD
E-CAD/DAPI
N-CAD
N-CAD/DAPI
Control
GPx2-OE
KRT14
KRT14/DAPI
VIM
VIM/DAPI
Control
GPx2-OE
D
KRT8
KRT14
KRT8/KRT14
PyMT2/Control tumors
Area 1
Area 2
Area 3
KRT8/KRT14
KRT8
KRT14
KRT8/KRT14
PyMT2/GPx2-OE tumors
Area 1
Area 2
Area 3
KRT8/KRT14
E
VIM
E-CAD
VIM/E-CAD
PyMT2/Control tumors
Area 1: E-CAD/VIM
Area 2: E-CAD/VIM
Area 3: E-CAD/VIM
VIM
E-CAD
VIM/E-CAD
PyMT2/GPx2-OE tumors
Area 1: E-CAD/VIM
Area 2: E-CAD/VIM
Area 3: E-CAD/VIM

**Fig. S5. Exogenous expression of GPx2 in PyMT2 cells inhibits EMT continuum.**

(**A**) Phase-contrast images of PyMT2 cells and PyMT2/GPx2-OE cells at 10X magnification showing morphological difference between the two cell lines. (**B**) Western blots showing protein levels of GPx2, N-CAD, SLUG, SNAI1, relative to ACTIN, in PyMT2 cells and PyMT2/GPx2-OE cells. (**C**) Immunofluorescent staining for E-CAD, N-CAD, KRT14, and VIM in PyMT2 control tumors (n=6) and PyMT2/GPx2-OE tumors (n=6) from 3 mice each. Three random sections from each tumor were immunostained. Representative images are shown. (**D-E**) Co-immunostaining of KRT8 with KRT14 (D), or VIM with E-CAD (E) in PyMT2 control tumors and PyMT2/GPx2-OE tumors from 3 mice each.

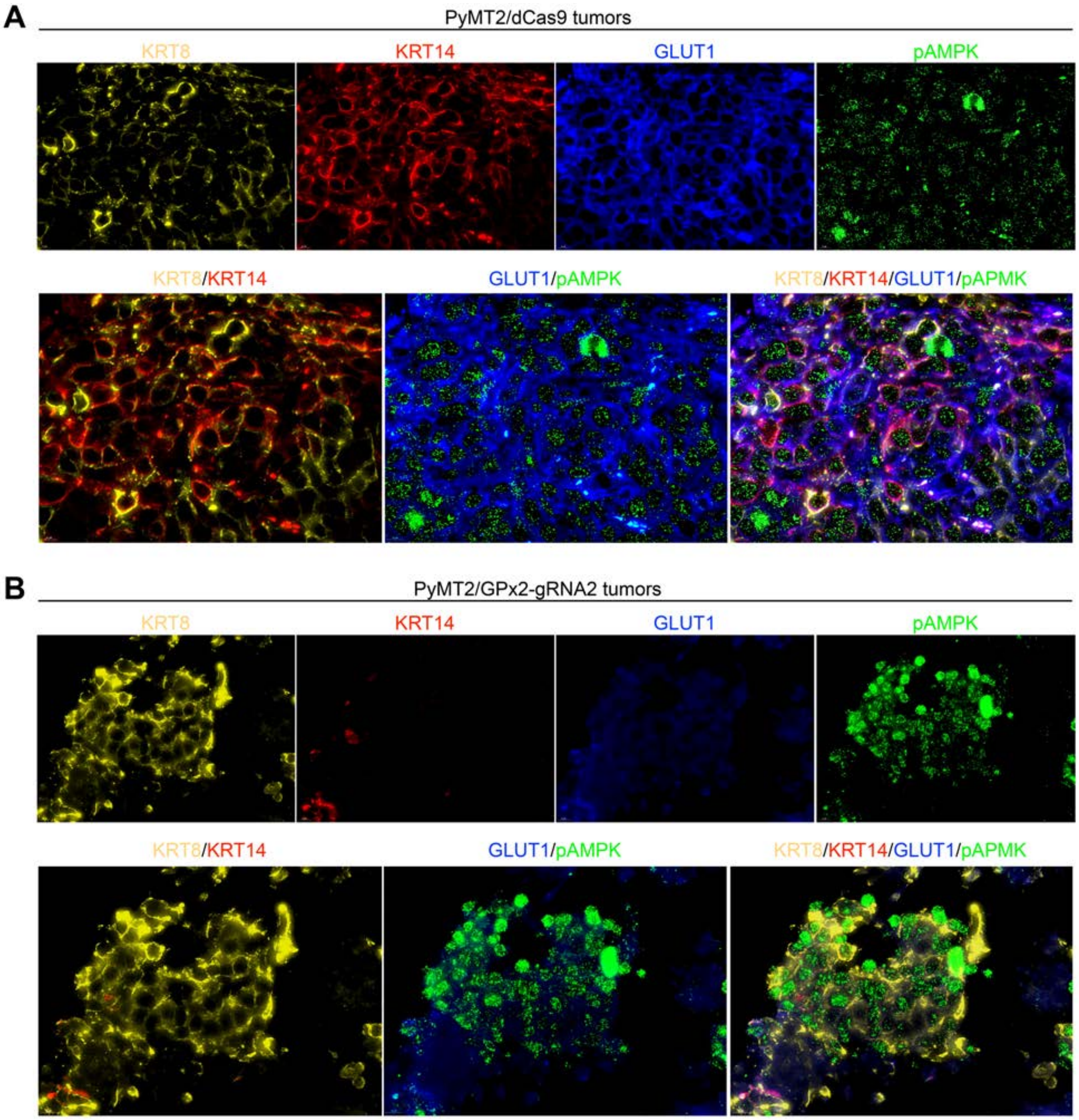


**Figure S6. Endogenous induction of GPx2 in metastatic PyMT2 cells attenuates the E/M and OXPHOS/Glycolysis hybrid phenotypes.** (**A-B**) Co-staining for KRT8, KRT14, GLUT1 pAMPK in PyMT2/dCas9 control versus PyMT2/GPx2-gRNA2 tumors (n=6) from 3 mice each.
